# Sequence-Dependent Dynamics of Synthetic and Endogenous RSSs in V(D)J Recombination

**DOI:** 10.1101/791954

**Authors:** Soichi Hirokawa, Griffin Chure, Nathan M. Belliveau, Geoffrey A. Lovely, Michael Anaya, David G. Schatz, David Baltimore, Rob Phillips

**Author notes:** S.H., G.A.L., D.G.S., D.B., R.P. designed research. S.H. performed research. S.H., N.M.B., G.A.L., M.A., D.B., R.P. provided new reagents and analytical tools. S.H., G.C., N.M.B., G.A.L., D.G.S., D.B., R.P. analyzed data. S.H., G.C., N.M.B., G.A.L., D.G.S., D.B., R.P. wrote paper. The authors declare no conflict of interest.

## Abstract

Developing lymphocytes in the immune system of jawed vertebrates assemble antigen-receptor genes by undergoing large-scale reorganization of spatially separated V, D, and J gene segments through a process known as V(D)J recombination. The RAG protein initiates this process by binding and cutting recombination signal sequences (RSSs) composed of conserved heptamer and nonamer sequences flanking less well-conserved 12- or 23-bp spacers. Little quantitative information is known about the contributions of individual RSS positions over the course of the RAG-RSS interaction. We employ a single-molecule method known as tethered particle motion to quantify the formation, stability, and cleavage of the RAG-12RSS-23RSS paired complex (PC) for numerous synthetic and endogenous 12RSSs. We thoroughly investigate the sequence space around a RSS by making 40 different single-bp changes and characterizing the reaction dynamics. We reveal that single-bp changes affect RAG function based on their position: loss of cleavage function (first three positions of the heptamer); reduced propensity for forming the PC (the nonamer and last four bp of the heptamer); or variable effects on PC formation (spacer). We find that the rare usage of some endogenous gene segments can be mapped directly to their adjacent 12RSSs to which RAG binds weakly. The 12RSS, however, cannot explain the high-frequency usage of other gene segments. Finally, we find that RSS nicking, while not required for PC formation, substantially stabilizes the PC. Our findings provide detailed insights into the contribution of individual RSS positions to steps of the RAG-RSS reaction that previously have been difficult to assess quantitatively.

**Summary:** V(D)J recombination is a genomic cut-and-paste process for generating diverse antigen-receptor repertoires. The RAG enzyme brings separate gene segments together by binding the neighboring sequences called RSSs, forming a paired complex (PC) before cutting the DNA. There are limited quantitative studies of the sequence-dependent dynamics of the crucial inter-mediate steps of PC formation and cleavage. Here, we quantify individual RAG-DNA dynamics for various RSSs. While RSSs of frequently-used segments do not comparatively enhance PC formation or cleavage, the rare use of some segments can be explained by their neighboring RSSs crippling PC formation and/or cleavage. Furthermore, PC lifetimes reveal DNA-nicking is not required for forming the PC, but PCs with nicks are more stable.

---

Jawed vertebrates call upon developing lymphocytes to undergo a genomic cut-and-paste process known as V(D)J recombination, where disparate gene segments that do not individually code for a protein are systematically combined to assemble a complete, antigen receptor-encoding gene (1). V(D)J recombination supports the production of a vast repertoire of antibodies and T-cell receptors that protect the host organism from a broad array of pathogens. However, gene segment combinations are not made in equal proportions; some gene segment combinations are produced more frequently than others (2–5). Although V(D)J recombination requires careful orchestration of many enzymatic and regulatory processes to ensure functional antigen receptor genes whose products do not harm the host, we strip away these factors and focus on the initial stages of V(D)J recombination. Specifically, we investigate how the dynamics between the enzyme that carries out the cutting process and its corresponding DNA-binding sites adjacent to the gene segments influence the initial stages of recombination for an array of synthetic and endogenous binding site sequences.

The process of V(D)J recombination (schematized in Fig. 1) is initiated with the interaction between the recombination-activating gene (RAG) protein complex and two short sequences of DNA neighboring the gene segments, one that is 28 bp and another that is 39 bp in length. These recombination signal sequences (RSSs) are composed of a well-conserved heptamer region immediately adjacent to the gene segment, a more variable 12- (for the 12RSS) or 23-bp (for the 23RSS) spacer sequence and a well-conserved nonamer region. For gene rearrangement to begin, RAG must bind to both the 12- and the 23RSS to form the paired complex (PC) state (Fig. 1B). Throughout the binding interaction between RAG and either RSS, RAG has an opportunity to nick the DNA (Fig. 1B zoom in) (6). RAG must nick both RSSs before it cleaves the DNA adjacent to the heptamers to expose the gene segments and to create DNA hairpin ends (Fig. 1C). DNA repair proteins complete the reaction by joining the gene segments to each other and the RSSs to one another (Fig. 1D).

**Fig. 1.**
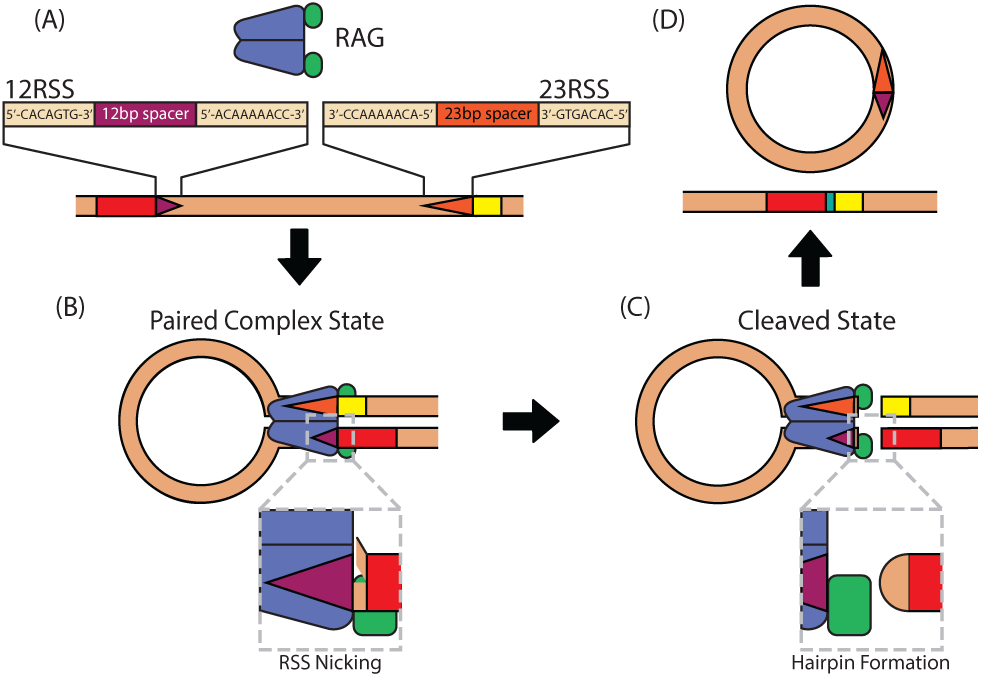
Schematic focusing on the initial steps of V(D)J recombination. (A) The RAG protein complex binds to the 12- and 23RSSs (purple and orange triangles, respectively) neighboring gene segments (shown as red and yellow boxes on the DNA), (B) forming the paired complex (PC). At any point when it is bound to an RSS, RAG can introduce a nick in the DNA between the heptamer and gene segment (shown with the magnified 12RSS) and must do so to both sites before (C) it cleaves the DNA to expose the gene segments. As indicated by the magnified gene segment end, the exposed DNA strands of the gene segment are connected to form a DNA hairpin. (D) Additional proteins join these segments together. In this work, the stages subsequent to DNA cleavage are not monitored.

RSS sequence-conservation studies across many organisms have shown a vast diversity of 12- and 23RSS sequences, mainly found through heterogeneity in the spacer region (7). Bulk assays reveal that changing an RSS sequence can significantly influence the RAG-RSS interaction and ultimately the success rate of completing recombination (8–12). Recent structural results provide evidence that RAG binding is sensitive to base-specific contacts and the local flexibility or rigidity of the 12- and 23RSS (13–15). Despite this extensive characterization on the interaction, little is known about how a given RSS sequence affects each step of the RAG-RSS reaction. In this work, we provide one of the most comprehensive studies of how RSS sequences govern the initial steps of V(D)J recombination and provide a quantitative measure of their effects on the formation frequency, lifetime, and cleavage probability of the PC.

We employ a single-molecule technique known as tethered particle motion (TPM) in which an engineered strand of DNA containing a 12RSS and 23RSS is attached to a glass coverslip at one end and to a polystyrene bead at the other (Fig. 2A). Using brightfield microscopy, we collect the root mean squared displacement (RMSD) of the bead over time to identify the state of the RAG-RSS interaction. As illustrated in Fig. 2B, when RAG forms the PC with the RSSs, the DNA tether is shortened, constraining the motion of the bead which is manifest in a reduction of the RMSD. When RAG cleaves the PC, the bead is released and diffuses away from the tether site (Fig. 2C). TPM has been applied to track the dynamic behavior of various protein-DNA systems, including RAG and RSS (16–21). It is with the temporal resolution provided by TPM that we can track the full progression of individual RAG-RSS interactions from PC formation to cleavage.

**Fig. 2.**
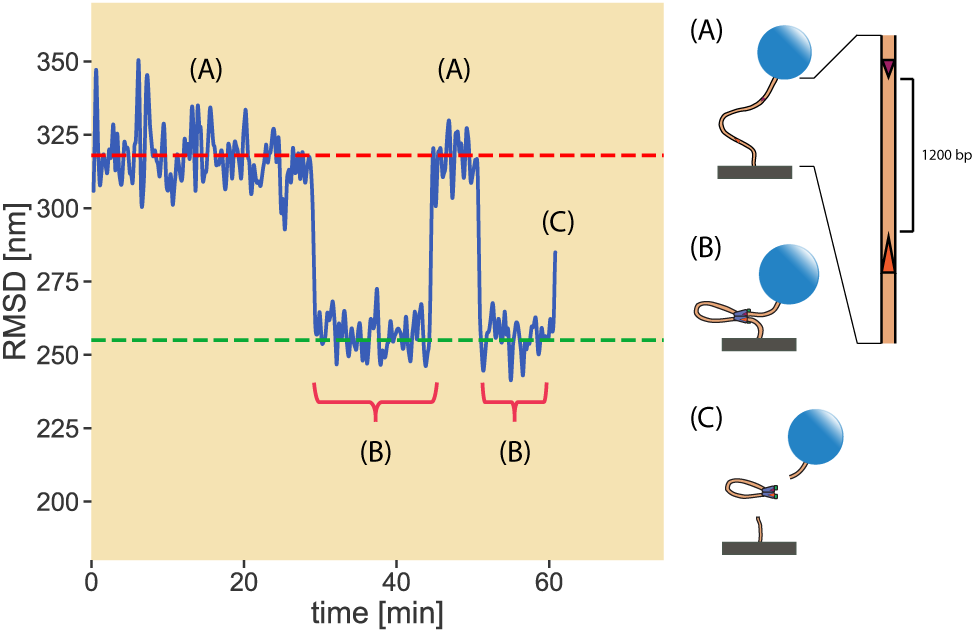
Sample data output of TPM. By tracking the root mean square displacement (RMSD) of the tethered bead position undergoing restrained Brownian motion, we discern when the DNA tether is (A) in the unlooped state, (B) in the PC, or looped, state and (C) cleaved. Red brace shows the measured PC lifetime. The dashed horizontal lines distinguish the unlooped (red) and looped (green) states of the DNA, and are drawn before examining the bead trajectories and based on the length of the DNA tether and the distance between the RSSs along the strand, the extent to which HMGB1, a protein that binds nonspecifically to DNA and helps facilitate RAG binding, kinks the DNA and a set of calibration experiments relating the range of motion of the bead to the length of its tether. As depicted in the zoom-in on the DNA in (A), the distance between the 12RSS and 23RSS is fixed at 1200 bp.

We were interested in using TPM to determine the extent to which endogenous RSSs dictate the usage frequency of their neighboring gene segments and, for those RSS positions that do seem to influence gene segment usage, identify the steps in the RAG-RSS reaction when the RSSs help or hurt their gene segment. We first examine single bp changes to a designated reference RSS, thereby establishing a mechanistic understanding of the contribution of individual nucleotide positions to RAG-RSS dynamics. With the synthetic RSSs providing context, we study a set of endogenous RSSs, each of whose sequences can be directly related to the reference sequence and a subset of the characterized synthetic RSSs. This selection of RSSs was also chosen from repertoires where the usage frequencies of their gene segments are known. Finally, we report on why our attempts to uncover deeper quantitative insights on the kinetics of the system from the PC lifetimes were met with strong disagreement between our intuited model and the TPM data, and what that consequently says about our understanding of the molecular details of the RAG-RSS reaction. As this study resulted in a wealth of data on a large number of RSS sequences, we have developed an interactive online resource for visualizing the dataset in its entirety (22).

## Results

### Synthetic RSSs

We chose a 12RSS flanking the immunoglobulin *κ* variable (*Igκ*V) gene segment, V4-57-1, as the reference sequence due to its use in a previous TPM study on RAG-RSS interactions (20) for our reference sequence. This sequence has also been used in structural studies of RAG-RSS complexes (13, 15), allowing us to compare our results with known information on the RAG-RSS structure. To explore how RAG-RSS interactions are affected by single bp changes, we examined 40 synthetic RSSs consisting of single bp changes across 21 positions of the V4-57-1 12RSS, with a particular focus on altering the spacer which is the least well-understood element in the RSS. We also studied changes made to positions 3-7 of the heptamer and various positions of the nonamer. The first three positions of the heptamer are perfectly conserved (7), likely to support DNA distortions needed for nicking and for base-specific interactions with the cleavage domain on RAG1 after nicking (13–15), while heptamer positions 4-7 also mediate base-specific interactions with RAG (13). The nonamer is bound by a nonamer-specific binding domain on RAG (13, 23). Throughout our synthetic and endogenous RSS study, we used the same concentrations of the purified forms of the two proteins that make up RAG (RAG1 and RAG2) and the high mobility group box 1 (HMGB1) protein, which binds nonspecifically to DNA and helps facilitate RAG binding to the RSSs (12). We also fixed the distance between the two binding sites to be 1200 bp, thereby constraining our study to the influence of binding site sequence on RAG-RSS dynamics alone. In addition, all of these endogenous 12RSSs are partnered with a well-characterized 23RSS (13, 15, 20) adjacent to the frequently-used gene segment from the mouse V*κ* locus on chromosome 6 (5). The sequence of this RSS is provided in Table S1 of the SI text.

We pooled the relevant data across experimental replicates to characterize synthetic RSSs by three empirical properties, namely the frequency of entering the PC (looping frequency), the quartiles of the PC lifetime (dwell time) distribution, and the probability of exiting the PC through DNA cleavage (cutting probability). We define the looping frequency as the ratio of distinct PCs observed to the total number of beads monitored over the course of the experiment. Because a single DNA tether can loop and unloop multiple times over the course of the experiment, the looping frequency can in principle range from 0 to *∞*. The dwell times were obtained from measuring the lifetimes of each PC state, irrespective of whether the PC led to a cleavage event or simply reverted to an unlooped state. To compute the cutting probability, we considered the fate of each PC as a Bernoulli trial with cleavage probability *pcut*. The cutting probability reported here is the ratio of observed cleavage events to total number of observed PCs. A more detailed discussion of these calculations and the corresponding error estimates are provided in Materials & Methods and SI Text. We provide a detailed record of the data for the synthetic RSSs on the website. This webpage includes heatmaps to qualitatively illustrate how the synthetic RSSs differ in the three defined metrics. By clicking on a particular cell in any of the heatmaps, the interactive displays the measured looping frequency of the synthetic RSS with the corresponding bp change with several confidence intervals. In addition, the webpage shows for the RSS empirical cumulative distribution functions (ECDFs) of PC lifetimes in three groups: PCs that are cleaved, PCs that are unlooped, and both together. Finally, this webpage includes the complete posterior probability distribution of the cleavage probability for each synthetic RSS.

Fig. 3 illustrates the significant effect that a single bp change to an RSS can have on the formation (A), stability (B), and cleavage (C) of the PC, reaffirming that RSS sequence plays a role in regulating the initial steps of recombination. Of interest is the observed difference in phenomena between changes made to the third position and those made to the last four bases of the heptamer region. Bulk assays showing that deviating from the consensus C at heptamer position 3 essentially eliminates recombination (8, 10), yet we found that changes to G or T did not inhibit PC formation (Fig. 3B). In fact, these alterations showed similar looping frequencies and PC lifetimes (Fig. 3B) as found for the reference sequence. However, both synthetic RSSs almost completely suppress cleavage (Fig. 3C). We provide the full probability distribution for the estimate of the cutting probability (this posterior distribution is fully defined in the Materials & Methods and SI text) for these two RSSs in Fig. 3D. Nearly all of the distribution is concentrated below 10%, showing that cutting the PC is exceedingly rare. Thus, having a C at the third position of the heptamer is critically important for the cleavage step but is not necessary for RAG to form the PC with the RSS. Although deviations from the consensus nucleotide at the heptamer position 3 do not prevent RAG from forming the PC, they do impede DNA cleavage.

**Fig. 3.**
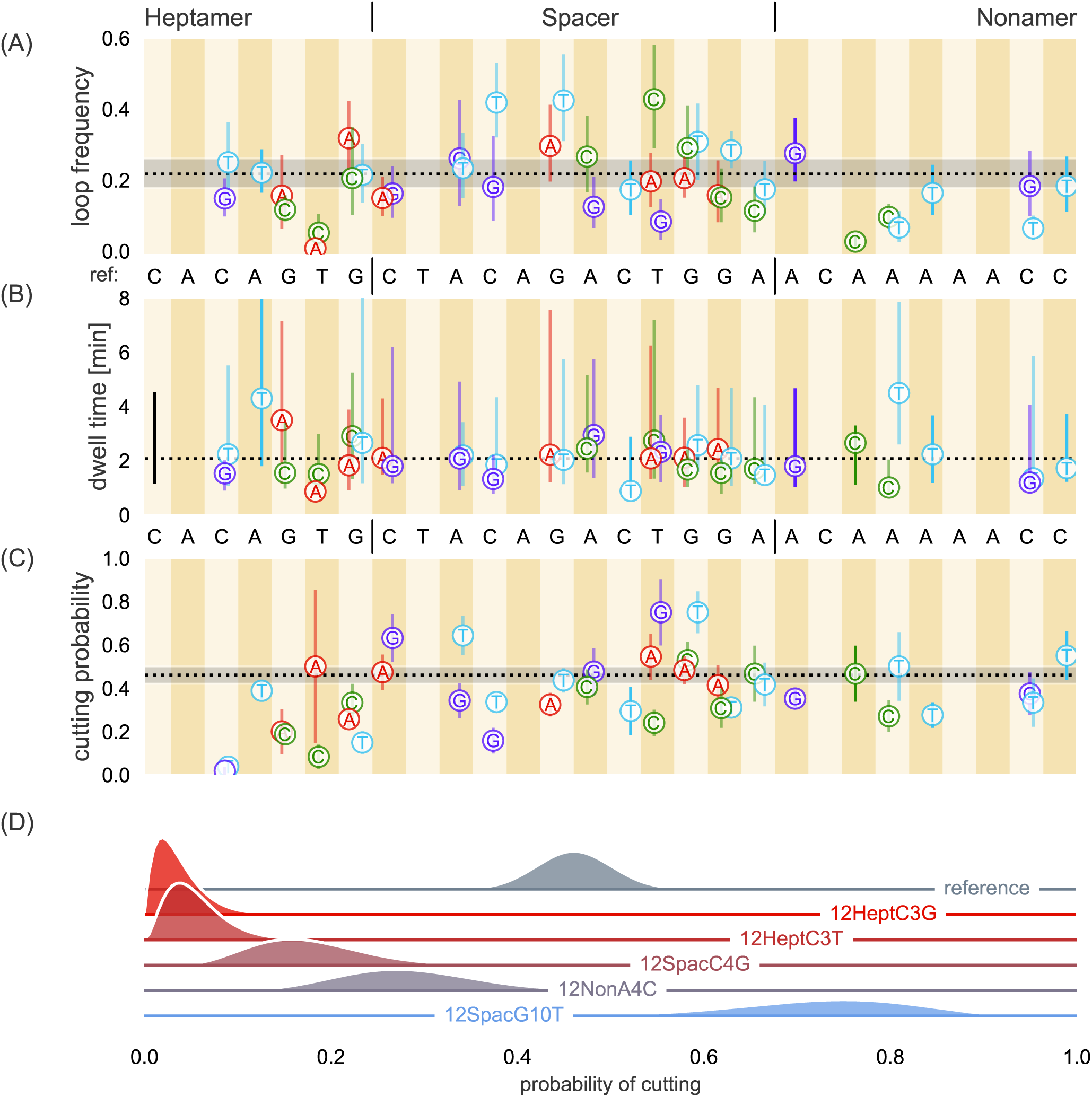
TPM data for single bp changes introduced at various positions of the reference 12RSS. (A) Loop frequency with 95% confidence interval, (B) dwell time with median represented as a point and lines extending to the first and third quartiles of the distribution and (C) cutting probability with standard deviations. The dotted black line in (A) is set at the reference loop frequency, 0.23, with shaded area denoting the extent of the 95% confidence interval for the reference. The dotted black line in (B) denotes the reference 12RSS median dwell time, 1.8 minutes, with the black bar at the left denoting the first and third quartiles of the distribution. Dotted black line in (C) is the most probable cutting probability for the reference sequence, roughly 0.4, with the grey shaded region setting one standard deviation. The reference sequence is provided along the x-axis for ease of determining the position where the change was made and the original nucleotide. The introduced nucleotide is provided in the figure with the letter and color-coded (red for A, green for C, light blue for T and purple for G). Heptamer, spacer, and nonamer regions are also separated by vertical lines in the sequences. (D) Ridgeline plot of posterior distributions of the cutting probability, given the number of loops observed and loops that cut (see SI) for a subset of the synthetic RSSs (labeled and colored along the zero-line of the respective ridgeline plot). Height of the distribution is proportional to the probability of a given cutting probability.

We find that PC formation is reduced compared to the reference sequence when the last four bases of the heptamer are altered, particularly at the fifth and sixth positions. Of more than 400 DNA tethers with the 12RSS containing a T-to-A change at heptamer position 6, we observed the PC only once, which subsequently led to cleavage. This result is consistent with recent findings that the consensus TG dinucleotide at the last two positions of the heptamer supports a kink in the DNA and may be critical for RAG binding (14). We notice that some changes increase the median time spent in the PC such as with the heptamer position 4 (Fig. 3B). This RSS also had one of the widest dwell time distributions of all of the synthetic RSSs studied. Furthermore, we find that the cutting probability decreased when we altered any one of the last four positions of the heptamer, but to a lesser extent than for changes made to the heptamer 3. The single bp change that had the greatest effect, located at heptamer position 6 (T to C) showed 2 out of 28 PCs led to cleavage.

Although we observed only modest differences in the median dwell times when we altered the reference sequence in the spacer region, some alterations substantially affected the looping frequency and cutting probability. The C-to-T change at spacer position 4 doubled the frequency of observing the PC while a T-to-G change at the ninth position reduced PC formation nearly as much as changes made at heptamer position 6. These two changes made to the spacer reflect the observed extremes of spacer sequence effects on the looping frequency. While many of the changes in the spacer region do not alter the cutting probability, we can still find spacer-altered RSSs that improve or inhibit cleavage in this region. Fig. 3D shows that changing the fourth position from C to G reduces the cleavage probability, while altering the tenth position of the spacer from G to T increases the cleavage probability as well as the frequency of PC formation (Fig. 3A). RAG1 makes contacts along the entire length of the 12RSS spacer (14), helping to explain our finding that changes to the spacer can substantially alter the probability of PC formation and cutting, thereby playing more of a role than simply separating the heptamer and nonamer sequences.

Similar to spacer changes, most nonamer changes alter the PC dwell time by ≈1 minute relative to the reference sequence. However, unlike spacer-modified RSSs, most synthetic-nonamer RSSs reduced the frequency of PC formation. Disruptions to the poly-A sequence in the center of the nonamer cause a substantial reduction in looping frequency, most notably the near complete inhibition of PC formation with the A-to-C change at nonamer position 3. This detrimental effect of deviating from the poly-A tract agrees with previous work demonstrating numerous protein-DNA interactions in this region and with the proposal that the rigidity produced from the string of A nucleotides is a critical feature for RAG1 to bind the nonamer (14, 23). Furthermore, this reduction in looping frequency extends to changes made at the eighth and ninth positions. The sequence deviations in the nonamer region, however, do not significantly affect cleavage once the PC has formed, as evidenced by the overlap in the posterior distributions of the reference sequence and its nonamer variant with the greatest reduction in the cleavage probability (position 4, A to C), in Fig. 3D. Overall, synthetic-nonamer RSSs have negative effects on PC formation with minimal effects on subsequent DNA cleavage, consistent with extensive biochemical and structural evidence that the primary function of the nonamer is in facilitating RAG-DNA binding (23).

### Endogenous RSSs

To build on our study of single bp effects on RAG-RSS dynamics, we selected a set of endogenous RSSs based on existing gene usage frequency data and whether their sequences were superpositions of the reference RSS and subsets of our studied synthetic RSSs. Specifically, we selected gene segments from the mouse V*κ* locus on chromosome 6 from data collected by Aoki-Ota *et al.*, including a variety of frequently-used gene segments (V1-135, V9-120, V10-96, V19-93, V6-15 and V6-17), two moderately-used gene segments (V4-55 and V5-43) and two rarely-used (V4-57-1 and V8-18) gene segments (5). The V4-57-1 12RSS is identical to the reference 12RSS from our synthetic RSS study. In addition, we examined DFL16.1, the most frequently used D gene segment from the murine immunoglobulin heavy chain (*Igh*) locus on chromosome 12 (4, 24). Unlike the V*κ* gene segments, which only need to combine with one gene segment, D gene segments must combine with two other gene segments to encode a complete protein. As a result, DFL16.1 is flanked on both its 5’ and 3’ sides by distinct 12RSS sequences, denoted DFL16.1-5’ and DFL16.1-3’, respectively, both of which are examined in this study. The sequences of all endogenous RSSs studied here are provided in the SI Text. We apply TPM on these sequences to determine whether their involvement in the RAG-RSS reaction could both provide insight into the usage frequency of their flanking gene segments and be predicted based on the activity profile of the synthetic RSSs.

To develop a better sense for how RAG interacts with these RSSs in their endogenous context, the 6 bp coding flank sequence adjacent to the heptamer of all but the V4-57-1 RSS was chosen to be the natural flank provided by the endogenous gene segment. RAG interacts with the coding flank during DNA binding and PC formation (13–15) and coding flank sequence can influence recombination efficiency, particularly the two bp immediately adjacent to the heptamer (25–27). Two T nucleotides and to some extent even a single T immediately 5’ of the heptamer inhibit the nicking step of cleavage and thus reduce recombination efficiency (25–27). We did not extensively analyze the contribution of coding flank sequence in this study, and only V6-15 RSS among the studied RSSs would be predicted to be detrimental due to the T flanking the heptamer; all other coding flanks have combinations of A and C as the two terminal coding flank bases. We kept the same coding flank for the V4-57-1 RSS as in a previous study (20) to facilitate closer comparison of the results of the synthetic RSSs. We do not expect much difference between the endogenous coding flank sequence (5’-CACT<u>CA</u>, where the two nucleotides closest to the heptamer are underlined) and the coding flank used here (5’-GTCG<u>AC</u>) because the two terminal coding flank bases are similar to those of all but the V6-15 RSS and for reasons discussed in the Discussion and SI Text. The coding flank sequences for all studied endogenous RSSs are included in the SI text. We present the results of the RAG-endogenous RSS interaction in Fig. 4 and provide an interactive tool for exploring these data on the paper website. This webpage includes an interactive feature where the looping frequency, ECDFs of looping lifetimes, and probability distribution of the cleavage probability of any two endogenous RSS can be directly compared.

**Fig. 4.**
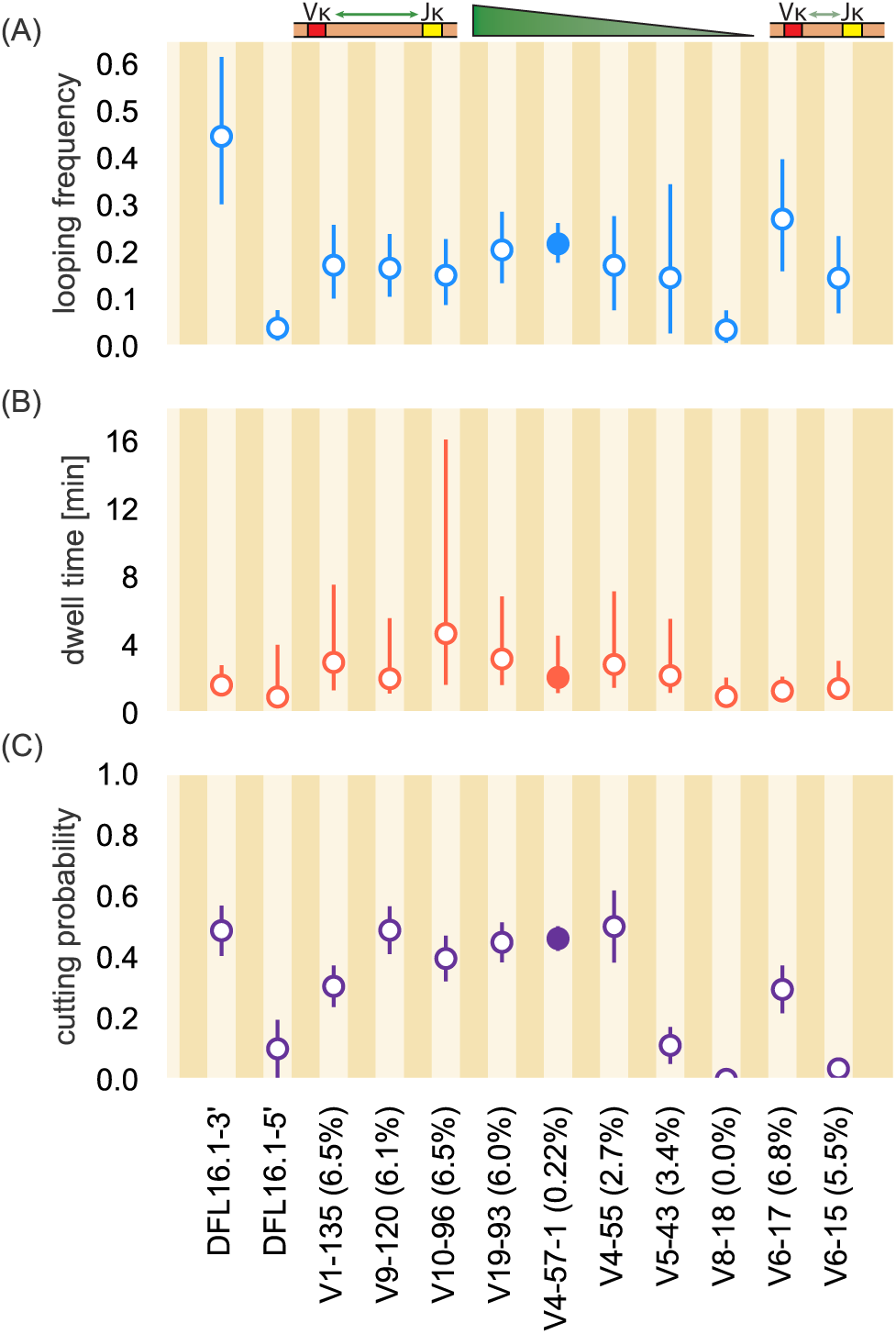
Observed dynamics between RAG and endogenous RSS sequences. (A) Frequency of PC formation (looping frequency) with 95% confidence interval. (B) Median PC lifetime with the lower error bar extending to the first quartile and the upper error bar extending to the third quartile. (C) Probability of DNA cleavage (cutting probability) of RAG with error bars showing one standard deviation. For discussion of the errors in Fig. 4A and 4C, see the SI text. DFL16.1-3’ and DFL16.1-5’ flank the same gene segment but in different orientations on the Igh chromosome. As shown in the graphic above Fig. 4A, V*κ* gene segments listed are ordered by their position along the chromosome, with linear distance from the J*κ* gene segments decreasing from left to right. Numbers in parentheses next to V*κ* gene segment denote percentage of usage in repertoire (5). The V4-57-1 12RSS has a filled in circle to denote that it is the reference sequence for examining the effects of single base changes.

The variable nature of all three metrics [looping frequency (Fig. 4A), dwell time (B), and cutting probability (C)] across RSSs highlights how, similar to the synthetic RSSs, endogenous sequences influence formation, stability, and cleavage of the PC differently. Of particular interest is the behavior of DFL16.1-3’ which shows the highest propensity for PC formation but some of the shortest PC lifetimes. Despite this short median dwell time, the probability of the PC successfully proceeding to DNA cleavage is approximately 0.5. Notably, the frequency of PC formation and the probability of cleavage are both greatly reduced for DFL16.1-5’ as compared to DFL16.1-3’, although their median PC dwell time and the width of the dwell time distributions are approximately equal. Reduced function of DFL16.1-5’ relative to DFL16.1-3’ is consistent with prior studies (24, 28, 29) and is addressed further in the Discussion.

The endogenous RSSs of the V*κ* gene segments show varying degrees of PC formation and cleavage probabilities. Many of the endogenous RSSs studied here, including those of gene segments used frequently *in vivo* (V1-135, V9-120, V10-96, V19-93, V6-17 and V6-15), demonstrate looping frequencies between 15 and 30 events per 100 beads. Gene segments V4-57-1 and V4-55 are used with low and modest frequency, respectively, yet in our experiments, they enter the PC with comparable frequency (approximately 20 to 30 loops per 100 beads). In general, we find these two sequences to behave almost identically in our experimental system, illustrating that other biological phenomena, such as higher-order DNA structure, govern the segment usage *in vivo* (4, 30). The endogenous V8-18 12RSS exhibits infrequent PC formation and cleavage and short median PC lifetimes, much like the DFL16.1-5’ 12RSS. Using the V8-18 12RSS, only 5 looping events were detected from 146 DNA tethers and cleavage was never observed. Despite the similarities in reaction parameters for the V8-18 and DFL16.1-5’ RSSs, DFL16.1 is the most frequently used D gene segment in the repertoire (4) while V8-18 is never used (5). One possible explanation for this difference is that the DFL16.1-5’ 12RSS does not participate in recombination until after its gene segment has undergone D-to-J recombination and has moved into the RAG-rich environment of the “recombination center.” This relocation is thought to facilitate RAG binding to the 5’ RSS of the committed D gene segment (31, 32). In contrast, V8-18, like other V*κ* gene segments, must be captured into the PC by RAG that has previously bound to a J*κ* RSS in the recombination center.

Fig. 4B demonstrates that, with the exception of the V10-96 RSS, PC lifetimes are similarly distributed across the endogenous RSSs examined in this work. Most RSSs have median dwell times between 1 to 3 minutes with the V8-18 12RSS displaying the shortest-lived median dwell time of roughly 40-50 seconds. While most endogenous RSSs here have a similar range between the first and third quartiles (see interactive figure on the paper website), the V10-96 12RSS distribution is noticeably wider, with the first quartile of the distribution being a longer lifetime than the median lifetime for most endogenous RSS distributions and the third quartile of this RSS extending out to over 19 minutes. These observations suggest a similar stability of the PC for all but the V10-96 RSS once RAG manages to bind simultaneously to both 12- and 23RSSs.

Fig 4C indicates that six endogenous RSS sequences from V1-135 to V4-55 have comparable cutting probabilities ranging from 0.4 to 0.5. Considering that the less-frequently used V4-57-1 and V4-55 gene segments have 12RSSs that show similar cutting probabilities and looping frequencies to the 12RSSs of more frequently-selected gene segments, other factors appear to prevent their selection. The low probability of cutting with the V6-15 12RSS is particularly noteworthy, with the low cutting probability of about 0.05 indicating that RAG tends to easily break the looped state rather than commit to cleavage. However, this low cutting probability might be attributed to the T in the coding flank immediately adjacent to the heptamer. Other features of the system must dictate the high-frequency usage of V6-15 *in vivo* (5).

### Kinetic Modeling of the PC Lifetime Distribution

Figs. 3B and 4B show that the vast majority of median looping lifetimes ranged between 1 to 3 minutes with rare exceptions, suggesting similar dwell time distributions for many of the RSS variants. However, many of these synthetic and endogenous RSSs have different probabilities of DNA cleavage, suggesting that at the very least the rate of cutting changes. As TPM has been used to extract kinetic parameters for various other protein-DNA systems (17, 18, 33, 34), we used the distributions of the PC lifetimes to estimate the rates of unlooping and cutting for each RSS and to attempt to discern a deeper connection between RSS sequence and fate of the PC. We developed a simple model in which a PC state can have two possible fates: either simple unlooping of the DNA tether or cleavage of the DNA by RAG. We characterized each of these outcomes as independent yet competing processes with rates *k*_unloop_ and *k*cut for unlooping and DNA cleavage, respectively. If the waiting time distribution *t*_unloop_ or *t*_cut_ for each process could be measured independently where only one of the two outcomes was permitted to occur, one would expect the probability densities of these waiting times given the appropriate rate to be single exponential distributions of the form

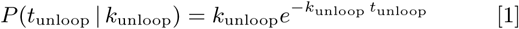

for the unlooping process and

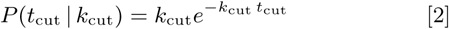

for DNA cleavage. However, as these two Poisson processes are competing, we cannot estimate *k*cut solely from the waiting time distribution of paired complex states that led to DNA cleavage nor *k*_unloop_ using the states which simply unlooped. As each individual cutting or unlooping event is assumed to be independent of all other cutting and unlooping events, the distribution of the dwell time *t* before the PC either unloops or undergoes cleavage can be modeled as an exponential distribution parameterized by the sum of the two rates,

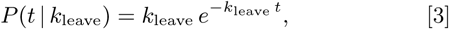

where *k*_leave_ = *k*_unloop_ + *k*_cut_.

Given the collection of waiting time distributions measured for each RSS, we estimated the values of *k*_leave_ which best describe the data. We find that the observed dwell times are not exponentially distributed for any 12RSS sequence analyzed, either endogenous or synthetic. Examples of these waiting time distributions along with an exponential distribution parameterized by the 95% credible region for *k*_leave_ can be seen for twelve of the RSS variants in Fig. 5. In general, the observed dwell times are underdispersed relative to a simple exponential distribution with an overabundance of short-lived PCs. We also find that the observed dwell time distributions are heavily tailed with exceptionally long dwell times occurring more frequently than expected for an exponential distribution. The ubiquity of this disagreement between our simple model and the observed data across all of the examined RSSs indicates that leaving the PC state either by reverting to the unlooped state or committing to the cleaved state is not a one-step process, suggesting that at least one of the two fates for the PC state on its own is not single-exponentially distributed as assumed in our null model of the dynamics.

**Fig. 5.**
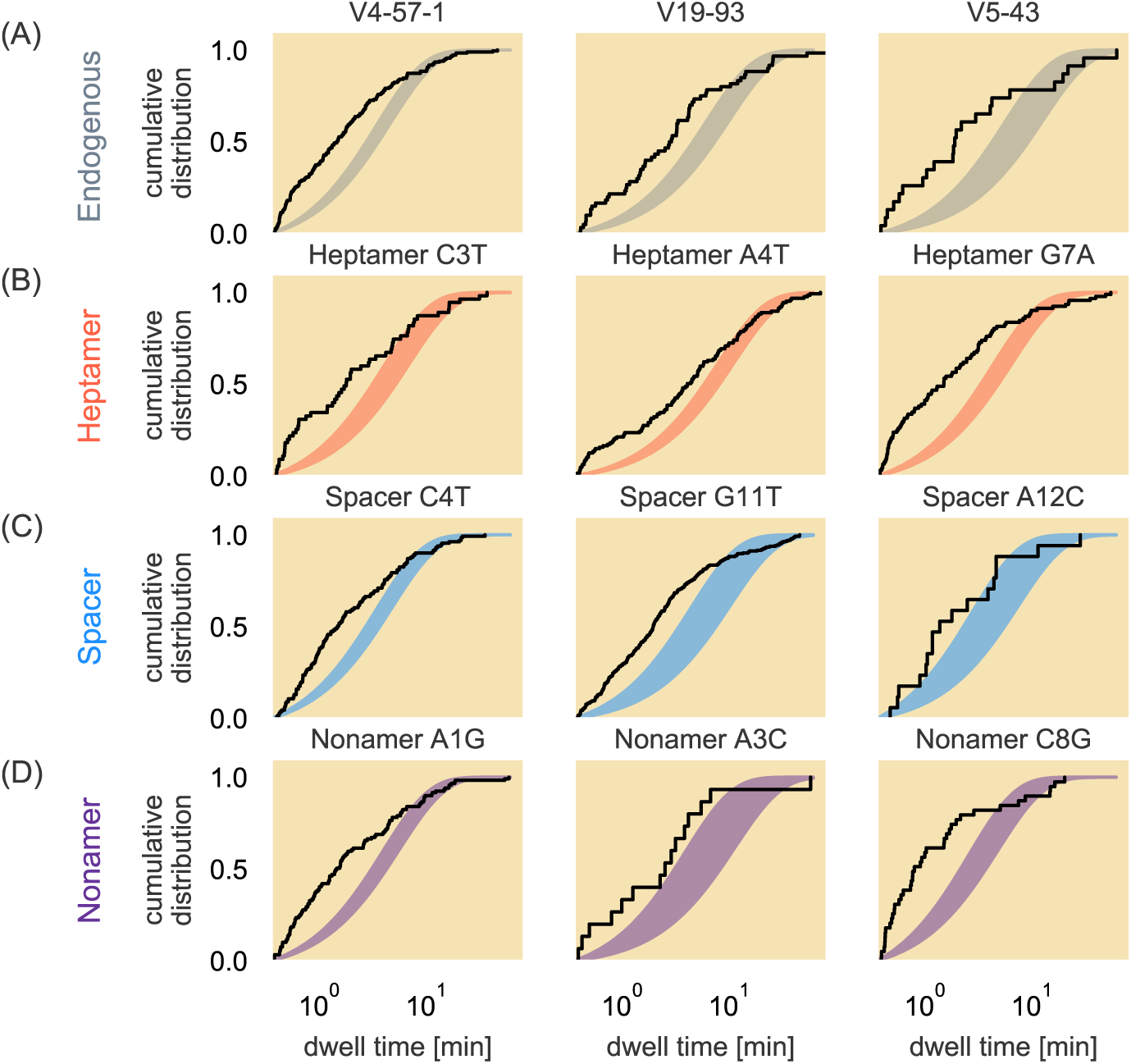
Non-exponential waiting time distributions for endogenous and synthetic 12RSSs. The empirical cumulative distribution of the measured PC lifetimes (black lines) are shown for representative endogenous sequences (A) as well as for the synthetic RSSs with single point alterations made in the heptamer (B), spacer (C), or nonamer (D) regions. The shaded area corresponds to the 95% credible region of a true exponential distribution parameterized in Eq. 3 given a posterior distribution for *k*_leave_, the rate of the arrival of either an unlooping or cleavage event.

One hypothesis for the disagreement between the model given in Eq. 3 and the data is that other processes, such as nicking of the DNA by RAG, create effects in the tethered bead trajectories that are too subtle to be detected in the TPM assays. Nicking creates a more stable RAG-single RSS complex (though this effect on PC stability had not been previously quantified) (13, 35) and can occur at any time after RAG binds to the RSS (6), making it exceedingly difficult to determine whether a given PC has one, both or neither of the RSSs nicked. As a result, we are unable to model the combined kinetics of unlooping and cleavage without also identifying when RAG nicks the RSSs to which it is bound.

Substitution of Ca^2+^ in place of Mg^2+^ in the reaction buffer allows RAG to bind the RSSs but blocks both nicking and cleavage (36), leaving unlooping as the only possible fate of a PC. To determine if unlooping could be modeled as a simple Poisson process, we measured the PC dwell time distribution for a subset of the RSSs in a reaction buffer containing Ca^2+^.

While we observe no cleavage of PCs in the Ca^2+^-based buffer, the dwell times of PC events are still not in agreement with an exponential distribution (left panels of Fig. 6A-C), indicating that the process of unlooping itself is not a Poisson process and that there are other looped states which our experimental system cannot detect. We also note that for each of the RSS variants the observed PC lifetimes are short lived compared to those in the Mg^2+^-based buffer, as can be seen in the bottom plots of Fig. 6. Because Ca^2+^ does not significantly alter DNA flexibility compared to Mg^2+^ (37), our data strongly argue that nicking itself results in a more stable PC. This is notable in light of recent structural evidence showing that nicking and the associated “flipping out” of two bases at the RSS-gene segment junction away from their complementary bases creates a more stable RAG-RSS binding conformation (13). With the more stable conformation from nicking one or both RSSs, the PC state can last longer than if RAG could not nick either RSS, which is reflected in the longer dwell time distributions when using Mg^2+^.

**Fig. 6.**
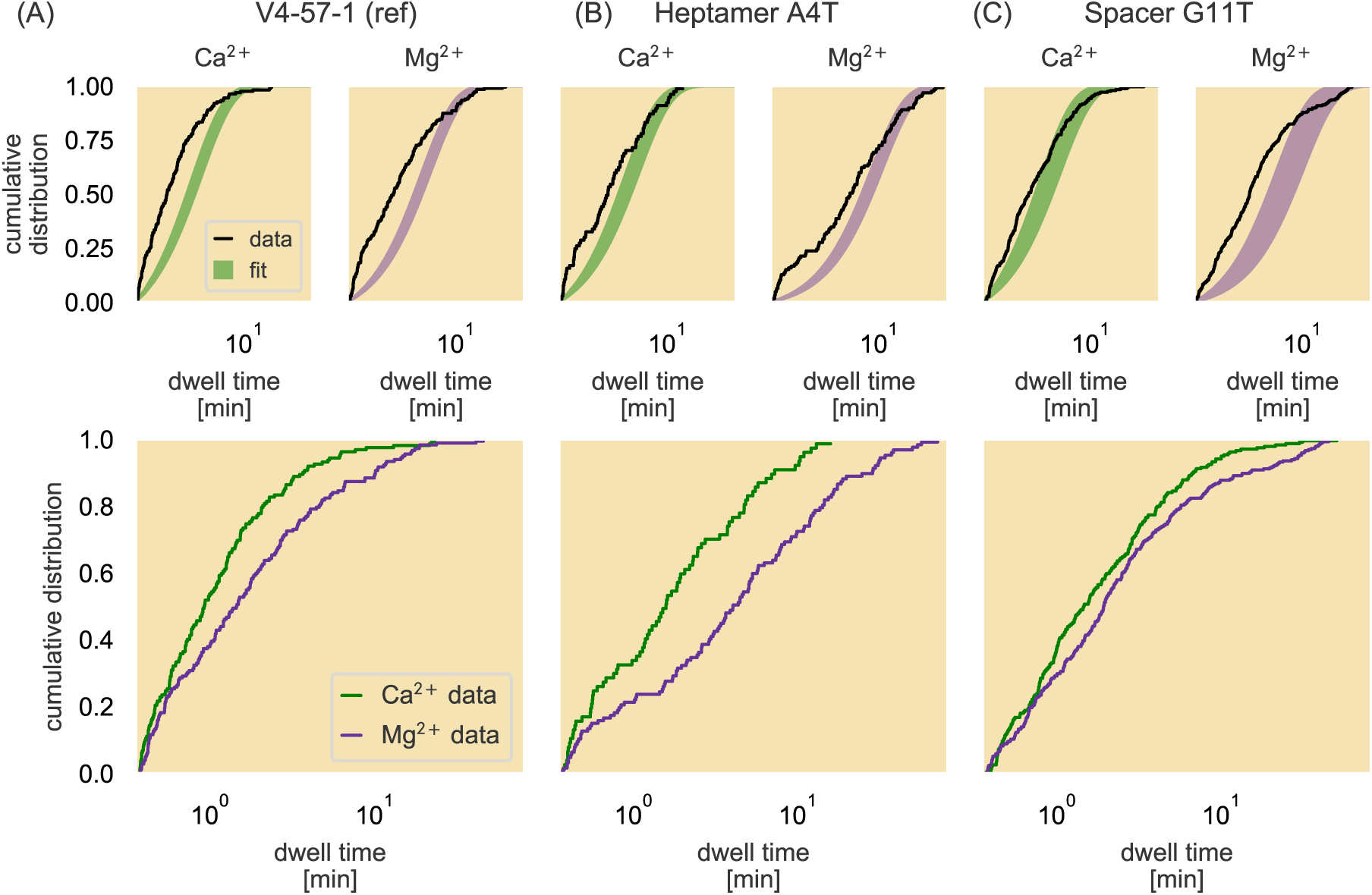
Empirical cumulative distributions of PC lifetimes with different divalent cations. The empirical cumulative dwell time distributions are plotted in black over the 95% credible region of the fit to an exponential distribution (top row) for the reference sequence (A), a base pair change in the heptamer region of the 12RSS (B), and a base pair change in the spacer region (C). The bottom plots show direct comparisons of the empirical cumulative dwell time distributions collected in either Ca^2+^- (green) or Mg^2+^- (purple) supplemented reaction buffer for each RSS.

## Discussion

Through the temporal resolution provided by TPM, we have discerned how RAG forms and cleaves the PC for a series of synthetic and endogenous RSSs. We find that the RSSs of frequently-used gene segments typically do not support more efficient PC formation or cleavage than those neighboring gene segments of more modest usage. This observation is consistent with recent findings that RSS strength, as assessed by the RSS information content (RIC) algorithm (11, 38–40), is only one of multiple parameters needed to be able to predict gene segment usage frequency (30, 41). Furthermore, we found from analyzing single bp variations of the V4-57-1 RSS that the efficiencies of PC formation and cleavage are sensitive to single bp changes depending upon the conservation level at the respective position. We see that altering the perfectly-conserved third position of the heptamer almost completely blocked cleavage by RAG, but did not significantly alter PC formation frequency or dwell time distribution. In contrast, most deviations from the consensus nucleotide at the last four positions of the heptamer or in the nonamer decreased the frequency of PC formation. Finally, even though few positions of the spacer have a consensus nucleotide (7), formation and cleavage of the PC can still be strongly affected by a single bp change in the spacer. In fact, sequence-context effects might help explain why some of these synthetic RSSs in less conserved positions of the spacer have such a strong influence on PC formation and cleavage on their own.

We asked to what extent we could account for the behavior of an endogenous RSS based on its constituent nucleotides as revealed by our synthetic RSS study. The comparative interactive tool on the paper website allows one to select an endogenous RSS to reveal not only its data on PC formation, PC lifetime distributions, and cleavage probability distributions, but also data for each nucleotide difference between it and the reference 12RSS through the relevant synthetic RSSs. Although we were unable to construct a quantitative model that could directly relate endogenous RSS behavior to the effects measured for each individual sequence deviation, these results provide several insights into the relation between RSS function and its constituent nucleotides. In particular, the data reveal a subset of RSS positions, including some in the spacer, that appear to strongly influence RAG-RSS interactions.

The synthetic RSS with the G-to-T change at the spacer position 10 strongly increases the cleavage probability and also enhances PC formation (Fig. 3A, C, D). These improvements might be due to the 5’-TG-3’ dinucleotide created by this change at spacer positions 10 and 11. Such a pyrimidine-purine (YR) pairing is inherently deformable (42) and a substantial 60*°*bend in the 12RSS is seen at this location in the spacer in RAG-RSS complexes (14). Hence, as noted previously (14), a YR combination at the 3’ end of the spacer in the 12RSS is favorable for DNA binding, consistent with our data. The DFL16.1-5’ RSS contains a T at spacer position 10 (Table S1), as well as several other nucleotides in the spacer that each individually increase PC formation (see the paper website), but this RSS hampers PC formation (Fig. 4A). Because spacer position 11 is also a T in the DFL16.1-5’ RSS, the T at position 10 does not create a YR pair and instead, the last seven bp of the spacer are all pyrimidines. A spacer with such a sequence might be particularly poor at supporting the DNA distortions needed for RAG-12RSS binding. This example of the importance of sequence context in determining how a particular bp will influence RSS function supports a concept borne out of the development of the RIC algorithm (11, 38, 40).

The contributions that coding flanks make to RAG-RSS dynamics (13) are important considerations to quantitatively model the RAG-DNA interactions, as each endogenous RSS neighbors a different coding flank. We attributed the low cleavage probability of the V6-15 RSS to the T immediately adjacent to the RSS in the coding flank, which has been shown to be detrimental to recombination efficiency (25–27). Because the other endogenous RSSs studied are rich in C and A nucleotides in the two bp adjacent to the heptamer, we compared data for two pairs of DNA constructs that differed only in coding flank sequence. One comparison involves the substrate containing the coding flank sequence used on the V4-57-1 RSS (5’-GTCG<u>AC</u>) and a substrate with a C-to-A change adjacent to the heptamer (5’-GTCG<u>AA</u>). The other pair is the V4-55 endogenous RSS substrate and the synthetic RSS substrate containing a C-to-A alteration at spacer position 1, where, fortuitously, the RSSs are identical and the coding flanks differ by five base pairs (5’-CACC<u>CA</u> for V4-55 and 5’-GTCG<u>AC</u> for the synthetic RSS). In both cases, the looping frequencies, PC lifetime distributions, and cutting probability distributions are similar for the respective pairs, arguing that these coding flank differences contribute little to the overall RAG-RSS reaction (see SI Text). Hence, coding flank differences present in all of the endogenous RSS substrates analyzed here, with the exception of the V6-15 RSS, are unlikely to have a strong influence on RAG-RSS dynamics. However, a more extensive examination of coding flank, particularly for G- and T-rich sequences, in a dynamic experimental method such as TPM will help to shed light on the extent to which these RSS-adjacent sequences influence the various steps of V(D)J recombination.

The V5-43 12RSS has a low level of PC formation, likely because of its C-to-T change at nonamer position 8, while its poor cutting probability can be attributed to a collection of sequence changes that reduce cleavage probability. The low frequency of PC formation with the V9-120 and V6-15 RSSs is likely driven primarily by the A-to-T change at nonamer position 4, with additional negative contributions coming from altering the reference spacer. And the DFL16.1-3’ RSS, which supported the highest frequency of PC formation across all RSSs studied, differ from the reference RSS at the fourth and sixth positions of the spacer that each in their own synthetic RSSs strongly stimulated PC formation. These findings support the important conclusion that spacer sequence can influence RSS synapsis by RAG.

We find that the DFL16.1-5’ RSS is much less competent or PC formation and cleavage than the DFL16.1-3’ RSS. Weaker activity of the 5’ RSS compared to the 3’ RSS is consistent with the results of recombination assays performed using plasmid substrates in cells (28, 29) and for chromosomal recombination when DFL16.1 was placed approximately 700 bp from its *Igh* J gene segment partner, J_H_1 (24). However, when assayed in their natural location over 50 kb from the J_H_ gene segments, the two RSSs support roughly equal levels of recombination as long as they are in the same orientation relative to the J_H_ 23RSSs (24). The existing data argue that the DFL16.1-5’ RSS is intrinsically less active for recombination than the DFL16.1-3’ RSS, but this difference can be minimized over large chromosomal distances when chromatin “scanning” by RAG is the dominant mechanism for bringing RSSs together to form the PC (24, 43).

Our study of both synthetic and endogenous RSSs explains the low usage of the V8-18 gene segment in the *Igκ* repertoire and further highlights the strong impact that can be exerted from a single nucleotide change to an RSS. The V8-18 RSS contributes to inefficient PC formation and further interrogating each sequence mismatch between the V8-18 and reference RSSs revealed that its T-to-A alteration at heptamer position 6 is sufficient to virtually abrogate PC formation. This result provides a mechanistic explanation for why the V*κ*A2b gene segment is underutilized in the antibody repertoire of Navajos, which in turn has been proposed to account for the high susceptibility of Navajos and several genetically-related groups of Native Americans to *Haemophilus influenza* type b infection (44). The V*κ*A2b RSS differs in sequence from the more common and efficiently recombined V*κ*A2a RSS by a single T-to-A change at the heptamer position 6 (44–46). We conclude that the inefficient recombination caused by this alteration is due to a defect in PC formation and suggest that any gene segment whose RSS contains an A at the sixth position of the heptamer will recombine poorly. Consistent with this, A is almost never observed at the sixth position of the heptamer in either the 12- or 23RSS (7).

Our attempts to obtain quantitative insight into the kinetics of RAG-RSS dynamics led to two interesting findings on the nature of the interaction. Upon first applying our fitting procedure to determine the rates of unbinding and cleavage, we learned that at least one of these two processes did not behave as a simple Poisson process. Thinking that our inability to detect nicking was the culprit, we examined the rate of unlooping in the absence of nicking by using Ca^2+^ instead of Mg^2+^ in our reactions. Here, our finding that the PC lifetimes were not exponential for any of the studied RSSs further thwarted our efforts to obtain a pure measurement of the rate of unlooping. These Ca^2+^ results suggest that the PC state may have multiple conformations like the *lac* repressor (47) in that the two RAG1/2 dimers may have multiple states, or that binding to the heptamer and to the nonamer on each RSS are actually separate sequential processes. One possible source of distinct conformations is the dramatic 180*°*rotation of the DNA that must occur prior to nicking. Rotated and unrotated configurations of un-nicked RSSs have been identified in recent structural studies (14, 15), but would be indistinguishable in the TPM assay. Despite these challenges to obtaining a quantitative description, our data demonstrate that nicking of an RSS is not a prerequisite for RAG to form the PC state, consistent with previous gel shift analyses performed either in Ca^2+^ or with RAG mutants lacking catalytic activity (48, 49). In addition, our findings demonstrate that PCs with nicked RSSs are more stable than those where RSSs cannot be nicked, extending previous findings made with RAG bound to single RSSs (35). To our knowledge, this study is the first attempt to obtain kinetic rates of unlooping from and cutting of the PC and reveals that there are still key details in the reaction that are left unaccounted for.

The work presented here leaves open several questions about the RAG-RSS dynamics. Although our TPM assay detects PC formation and cleavage, it does not detect nicking, preventing us from determining how the RSSs studied influence the rate of nicking or when nicking occurs relative to PC formation. Even without nicking, we see that the unlooping dynamics behave differently from a simple Poisson process. This result suggests a need for an experimental method such as single-molecule FRET (50) that can detect such subtle conformational changes that occur between RAG and the RSS. Finally, we have left the 23RSS unchanged in this study, but it is possible that the trends that we see for our synthetic or endogenous 12RSSs may change with a different partner RSS and shed more light on the “beyond 12/23 rule” (11, 51, 52). Ultimately, these finer details in the RAG-RSS interaction can provide a more complete kinetic description of the initial phases of V(D)J recombination. While we changed the 12RSS sequence in this work, the TPM assay in principle allows us to titrate other parameters, such as the distance between RSSs, or introduce more biochemical players to better contextualize our work in the bigger picture of recombination *in vivo*.

## Materials and Methods

### Protein purification

The two RAG components, core RAG1 and core RAG2 (RAG1/2), are purified together as outlined in (20). Maltose binding protein-tagged murine core RAG1/core RAG2 were co-expressed by transfection in HEK293-6E suspension cells in a 9:11 w/w ratio for 48 hours before purifying using amylose resin. HMGB1 is purified as outlined in (20). His-tagged HMGB1 was expressed in isopropyl-*β*-D-1-thiogalactopyranoside-induced BL21 cells for 4 hours at 30^°^C before purification. For more details, see the SI Text.

### Flow cell assembly

TPM flow cells were assembled by drilling four holes along each length of a glass slide before cleaning the slides and cover slips. The slides and cover slips were treated with an epoxidizing solution for at least an hour and a half. Upon completion of the treatment, flow cells are assembled by cutting four channels into double-sided tape to connect the drilled holes at opposite ends of the glass slide before adhering to the cover slip on one side and the glass slide on the other. Short connective tubes are inserted into each of the holes to serve as inputs and outputs for fluids and sealed using 5-minute epoxidizing solution. The constructed flow cells are baked for fifteen minutes on the hot plate.

### Tethered bead assembly

Tethered beads are constructed by incubating antdigoxigenin in the flow cell channels for two hours to allow for sticking to the glass surfaces. After washing away excess antidigoxigenin in a buffer solution containing Tris-HCl, KCl, MgCl_2_, DTT, EDTA, acetylated BSA and casein, engineered strands of 2900 bp-long DNA containing a 12RSS and a 23RSS located 1200 bp apart and tagged with digoxigenin on one end and biotin at the other end are injected into the flow cells to attach the digoxigenin end of the DNA to the anti-digoxigenin-scattered surfaces. After excess DNA is washed out, 490 nm streptavidin-coated polystyrene beads are added to the channels and incubated for no more than 3 minutes to bind the biotin-labeled end of the DNA. Excess beads are washed away and the TPM assembly buffer is replaced with a RAG reaction buffer containing Tris-HCl, KCl, glyercol, DTT, potassium acetate, MgCl_2_, DMSO and acetylated BSA. For Ca^2+^ studies, CaCl_2_ is used in place of MgCl_2_ in the RAG reaction buffer and in the same concentration. See SI Text for visual demonstration of TPM preparation.

### TPM experiment

TPM experiments involve the simultaneous acquisition of bead trajectories from two different channels on separate microscopes. One of the channels contains tethered DNA with a 12RSS and a 23RSS oriented toward each other (nonamer regions on both RSSs closest to each other). Properly tethered beads are filtered using various methods to ensure proper spacing from neighboring beads and that individual beads are tethered by a single strand of DNA. The trajectories of the selected beads are then examined in the absence of RAG and HMGB1 for ten minutes before flowing in 9.6 nM murine core RAG1/core RAG2 and 80nM full-length HMGB1 and acquiring bead trajectories for at least one hour. Additional information on bead selection criteria and identification of PCs are provided in the SI Text.

### Statistical inference

We used both Bayesian and Frequentist methods in this work to calculate parametric and nonparametric quantities, respectively. The PC formation frequencies were assigned confidence intervals via bootstrapping. Briefly, the observed beads and their reported PC formation counts were sampled with replacement to generate a simulated data set of the same length as the number of observations. The looping frequency was then calculated as the total loops formed among the generated dataset divided by the number of beads and the distribution was resampled again. This procedure was performed 10^6^ times and we report various percentiles of these bootstrap replicates, as shown both in the main text and on the paper website.

To compute the cleavage probability and PC leaving rate *k*leave, we used a Bayesian definition of probability and constructed a posterior distribution for each as is explicitly laid out in the SI Text. The displayed posterior distributions for the cleavage probability were generated by numerically evaluating the posterior distribution over a range of cleavage probabilities bounded from 0 to 1. The reported values for the cleavage probability and uncertainty were computed analytically and is derived in the SI text.

To estimate *k*_leave_ we again constructed a posterior distribution. Here, we chose an exponential form for the likelihood and assumed an inverse Gamma distribution as a prior on the leaving rate. This posterior was then sampled using Markov chain Monte Carlo as is implemented in the Stan probabilistic programming language. A more detailed derivation of the posterior distribution is provided in the SI Text. All models and code for this inference are available on the paper website.

### Data and code availability

All data and code are publicly available. Raw image files can be obtained upon request. Preprocessed image data can be downloaded from CaltechDATA research data repository under the DOI:10.22002/D1.1288. Processed data files, Matlab, and Python code used in this work can be downloaded either from the paper website or on the dedicated GitHub repository (DOI:10.5281/zenodo.346571).

## ACKNOWLEDGMENTS

We thank members of the David G. Schatz, David Baltimore, and Rob Phillips labs for useful discussions and Caltech’s Protein Expression Center for supplying resources and equipment for protein purification. We also thank Helen Beilinson, Justin Bois, Zev Bryant, Heun Jin Lee, Stephanie Johnson, Eddy Rubin, Charlie Starr, Yuhang Zhang, and Haojie Zhuang for discussions. This work was supported by R01GM085286 and 1R35 GM118043 Maximizing Investigators’ Research Award (MIRA) (to R.P.). S.H. was also supported by the Caltech Center for Environmental Microbial Interactions (CEMI) (R.P.), the Foundational Questions Institute (FQXI) (R.P.), and the Sackler Foundation (D.B.).

