## Supplementary Information for "Sequence-Dependent Dynamics of Synthetic and Endogenous RSSs in V(D)J Recombination"

### Supplementary Information for "Probing the Sequence-Dependent Dynamics of Synthetic and Endogenous RSSs in V(D)J Recombination"

Soichi Hirokawa[a], Griffin Chure[b], Nathan M. Belliveau[b,c], Geoffrey A. Lovely[d], Michael Anaya[b], David G. Schatz[e], David Baltimore[b], and Rob Phillips[b,f,1]

[a]Department of Applied Physics, California Institute of Technology, Pasadena, CA, USA
[b]Division of Biology and Biological Engineering, California Institute of Technology, Pasadena, CA, USA
[c]Present Address: Howard Hughes Medical Institute and Department of Biology, University of Washington, Seattle, WA, USA
[d]National Institute on Aging, National Institutes of Health, Baltimore, MD, USA
[e]Department of Immunobiology, Yale University School of Medicine, New Haven, CT, USA
[f]Department of Physics, California Institute of Technology, Pasadena, CA, USA

October 3, 2019

**S3 Posterior Distributions of the Endogenous Sequences** **11**

**S4 Coding Flank Contributions** **12**

**S5 $Ca^{2+}$-$Mg^{2+}$ Looping Frequency Comparisons** **14**

**S6 Endogenous RSS Sequences** **16**

**S7 Cloning a Different 12RSS in Plasmids** **16**

**S8 Synthetic 12RSS Primers** **16**

**S9 Protein Purification** **19**
S9.1 Murine core RAG1 and core RAG2 Co-Purification . . . . . . . . . . . . . . . . . . . . 19
S9.2 HMGB1 Purification . . . . . . . . . . . . . . . . . . . . . . . . . . . . . . . . . . . . . . 19

### S1 Experimental Methods

#### S1.1 Microscopy Components and Configuration

All TPM experiments were performed using two Olympus IX71 inverted microscopes with bright-field illumination. Experiments were run in parallel where one microscope imaged a flow cell containing DNA without any RSSs while the other microscope collected data on DNA strands containing the fixed 23RSS sequence and the studied 12RSS. Each microscope is outfitted with a 60x objective (Olympus) and a 1920-pixel×1200-pixel monochromatic camera with a global shutter (Basler acA1920-155um). The camera is configured in an in-house Matlab image acquisition script to acquire images at a frame-rate of 30 Hz. Each optical set-up is calibrated to relate DNA of lengths ranging from 300 bp to 3000 bp to the root mean squared distance of their tethered beads.

#### S1.2 TPM Preparation

A schematic of the tethered bead assembly process as discussed in the Materials & Methods of the manuscript is shown in Fig. S1. All buffers and assembly components are added to the flow cells by gravity flow. After anti-digoxigenin has coated the coverslip surface, flow cell chambers are washed twice with TPM assembly buffer containing 20 mM Tris-HCl (pH 8.0), 130 mM KCl, 2 mM $MgCl_2$, 0.1 mM EDTA, 0.1 mM DTT, 20 µg/mL acetylated bovine serum albumin (BSA), and 3 mg/mL casein. Once washed, DNA tethers are added and diluted in the TPM assembly buffer to a concentration of roughly 2.5 pM. The tethers are allowed to incubate within the cell for 15 minutes, allowing for the digoxigenin-functionalized ends of tethers to attach to anti-digoxigenin-coated coverslip. Unbound excess DNA is then removed from the flow cell and custom-ordered streptavidin-coated beads (Bangs Labs) are added to the flow cells, binding the DNA at the biotin ends, and left to incubate for three minutes before flushing excess beads from system. The prepared flow cell chamber is then equilibrated with RAG reaction buffer containing 25 mM Tris-HCl (pH 7.6), 75 mM KCl, 0.05% glyercol, 1 mM DTT, 30 mM potassium acetate, 2.5 mM $MgCl_2$, 5% DMSO and 100 µg/mL acetylated BSA for TPM experiments involving nicking or else the same buffer except with $CaCl_2$ in place of and at the same concentration as $MgCl_2$ for RAG-RSS interactions in the absence of DNA nicking.

#### S1.3 Image Processing

Image processing is performed on a field of view in the same manner established by Han *et al.* [1, 2]. After acquiring 60 images over two seconds, beads are identified by setting an intensity threshold before filtering over object sizes. Smaller regions of interest (ROIs) are drawn around each marker identified as a bead. After initial processing, an additional 120 images over four seconds are acquired and processed by determining intensity-weighted center of masses of beads. The radial root mean squared displacement (RMSD) of the bead position is then determined using the 120 images and compared to the calibration curve based on the expected length of the DNA. Beads are accepted if their RMS values correspond to DNA lengths within 100 bp of their actual lengths for the paired complex assays ($l_{DNA} \approx 2900$ bp). Beads are then further processed to examine their symmetry of motion. After the correlation matrix for the bead position over the 120 frames is obtained, the eigenvalues of the matrix are then obtained, which yield the lengths of the major and minor axes of the beads range of motion. If the square root of the ratios of the maximum eigenvalue over the minimum eigenvalue is greater than 1.1, then the asymmetry of the motion is considered to be due to the bead being tethered to multiple DNA strands and is therefore rejected. The remaining beads are kept for data acquisition.

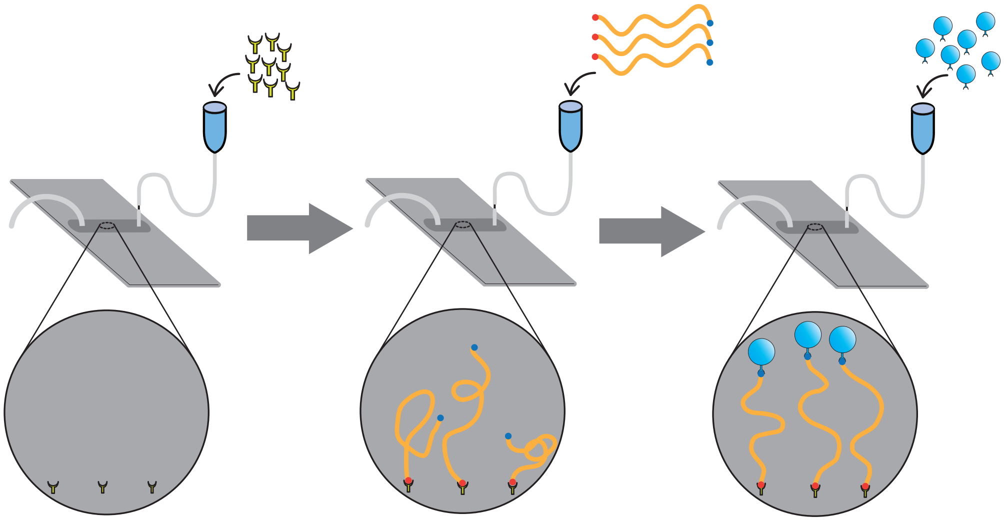

**Figure S1: Tethered bead preparation process.** Tethered beads are first assembled by adding anti-digoxigenin from Sigma-Aldrich into the flow cell chamber by gravity flow and left to incubate for at least two hours. The fluid is then displaced from the chamber by washing in TPM assembly buffer and introducing DNA tethers containing the desired 12RSS and a constant 23RSS. Unbound DNA tethers are then flushed out and streptavidin-coated beads are introduced to the flow cell. Once the tethered beads have been assembled, chambers are equilibrated with buffer used to study RAG-RSS reaction.

RMSD values of the bead center are obtained using a sliding window of 120 images acquired over four-second intervals. To correct for drift in the bead position, often due to the slow unidirectional motion of the microscope stage, the raw data are filtered through a first-order Butterworth filter with a cutoff frequency of 0.05 Hz. All ROI-binned image files can be downloaded from the CaltechDATA research repository under the DOI:XXXX. All code used to analyze these images can be found on the paper website or the paper GitHub repository (DOI: XXXX).

#### S2 Data Analysis: Extracting All Relevant Information from Bead Traces

All of the data reported and used in our results come solely from analyzing the RMSD as a function of time for each individual bead, hereafter called the "bead traces". This source must be further filtered in order to remove beads that passed through the initial image processing steps but still exhibit spurious behaviors, such as sticking to the glass surface or multiple beads falling into the same ROI and confounding the image processing. Information on the valid beads are then extracted and further analyzed through the bootstrapping method for the looping frequency confidence interval, the Bayesian analysis to obtain our posterior distributions of the cutting probability and the dwell time distributions for our analysis on kinetics of leaving the paired complex state.

##### S2.1 Selecting Beads for Further Analysis

Bead selection criteria after preprocessing is applied in the same manner as [1, 2, 3, 4]. After correcting for various systematic errors of the experiment, such as slow stage drift, beads are manually filtered based upon their RMSD trajectories both before and after introducing RAG and HMGB1. Tethers that have multiple beads attached are removed due to a larger variance in the RMSD trajectories for a given state. These beads can also be viewed through a software that shows the raw images at a defined time of the experiment. Furthermore, beads whose traces (in the absence of protein) lie below the expected RMSD value are considered to be a shorter DNA length than expected or an improperly tethered DNA strand and are also rejected. All other bead trajectories are tracked until one of four outcomes occurs: 1) RAG cleaves the DNA, causing a sharp increase in RMSD past the tether point and can be observed with the bead diffusing from the ROI; 2) the bead sticks to the glass slide for longer than a few minutes; 3) another bead enters the cropped region enclosing the studied bead due to stage drift that has not been correct or 4) the experiment ends, which typically runs for at least one hour of acquisition.

Once the beads have been selected, they are entered into an analysis that identifies whether a bead is in the unlooped or paired complex state using three thresholding RMSD values at every given instance of data acquisition, as performed in [2]. In instances where a bead trajectory drops below the minimum RMSD threshold, which is often an indication of temporary adhesion of the bead to the glass slide, or above the maximum RMSD threshold, set due to other temporary aberrations in bead motion, the time that the bead trace spent outside of these bounds are split evenly between the state that the bead was in immediately before and after. With the states of the bead defined at each time point, we can coarse-grain the bead trajectory into the amount of time spent in the paired complex or unlooped states. This allows us not only to determine the lifetime of each paired complex formed but also the number of loops that were formed for a given bead reporter. In addition, all looped states are assigned a binary number based on whether they subsequently led to unlooping (0) or to the bead untethering (1), the latter of which indicates DNA cleavage by RAG. Data on all beads kept by the TPM data acquisition code, including those that were manually filtered out during post-processing, are available on the CaltechDATA research data repository under the DOI:XXX.

#### S2.2 Bootstrapping Looping Frequency

While measuring the PC dwell time or the probability of PC cleavage is a straight-forward measurement, it is less clear how the propensity to enter the looped state should be calcuated. As described in the main text, we defined the looping frequency as the total number of observed PC events divided by the total number of beads observed over the experiment. It is tempting to simply repeat this calculation for each experimental replicate, average the results, and report a mean and standard error. However, the number of beads observed can vary greatly from one replicate to another. For example, one replicate may have 20 observed loop among 100 observed beads, bringing the looping frequency to 0.2. Another replicate of the same RSS may have 0 observed looping events, but among only 10 beads in total, bringing the looping frequency to 0. We would want to apply a penalty to the second measurement as we observed far fewer beads than in the first replicate, however assigning that penalty is also not obvious. To further complicate this calculation, some beads in an experiment will never undergo a looping event while others will show multiple events, making a bead-by-bead calculation of the looping frequency more challenging.

To address these challenges, we elect to compute and report the looping frequency as the total number of loops observed across all beads and experimental replicates, divided by the number of beads that were studied in total for that particular 12RSS. This metric, being bounded from 0 to $\infty$, accounts for the fact that for a given 12RSS, looping may occur many times. Furthermore, pooling the beads across replicates results in an appreciably large bead sample size, with the lowest sample size being greater than 80 beads and many RSSs having bead sample sizes in the hundreds.

In order to report a measure of the range of possible looping frequency values that could have been observed for a given RSS, we elect to apply bootstrapping. In bootstrapping as applied here, we treat the beads studied as the best representation of the population distribution of loop counts, as we do not have an idealized system where we could study infinitely many beads and track the number of paired complexes formed for each DNA tether. With this assumption that the experimentally-obtained loop count distribution provides the best representation of the population distribution, we can determine all possible ways we could have obtained the looping frequency by sampling from this empirical distribution. With this bootstrap-generated distribution of possible looping frequency values, we then calculate percentiles to provide confidence intervals on the looping frequency for comparison against the measured looping frequency. To see this in action, suppose our dataset on a particular RSS and salt condition contains $N$ tracked beads across all replicates, with bead $i$ reporting $l_i$ loops. Our measured looping frequency $f_{\text{meas}}$ would be $\frac{\sum_i l_i}{N}$. With bootstrapping, we can then determine our confidence interval on the measurement $f_{\text{meas}}$ given the bead dataset we obtained with TPM by following the general procedure:

1. Randomly draw $N$ different beads from the dataset of $N$ beads with replacement. This means that the same bead can be drawn multiple times.
2. Sum the total number of loops observed among this collection of $N$ beads and divide by $N$ to get a bootstrap replicate of the looping frequency, $f_{\text{bs},1}$.
3. Repeat this procedure many times. In our case, we obtain $10^6$ bootstrap replicates of the looping frequency.
4. For a confidence percentage $P$, determine the $(50 - \frac{P}{2})^{\text{th}}$ and $(50 + \frac{P}{2})^{\text{th}}$ percentiles from the generated list of $10^6$ bootstrapped calculations of the looping frequency.

As an example, we demonstrate this bootstrap method on the V4-57-1 12RSS, which we also refer to as the reference sequence for our synthetic RSS study. Through TPM, we had tracked 700 beads, each reporting some number of loops $l_i$. As a result, we draw 700 beads from this dataset with replacement in order to calculate a bootstrap replicate of the looping frequency. We repeat this $10^6$ times and obtain the result in Fig. S2. Although we report the 95% confidence interval in the manuscript, we also offer shades of the 5%, 10%, 25%, 50% and 75% confidence intervals on our website.

##### S2.3 Bayesian Analysis on Probability of Cuts

Bayesian analysis on cutting probability is applied in a similar manner to [5]. For a given RSS substrate ,to obtain the probability that RAG cuts a paired complex, $p_{cut}$, we construct a probability density function for $p_{cut}$ conditioned on our data. In this case, our data for each RSS examined is the total number of loops we observed in TPM, $N$, and the number of loops that were cut, $n$, so $n \leq N$. In short, we wish to determine the probability of $p_{cut}$ conditional on $N$ and $n$, or, written concisely, as $P(p_{cut}|N, n)$. Bayes' Theorem tells us that

$$P(p_{cut}|N, n)P(N, n) = P(n|N, p_{cut})P(N, p_{cut}). \tag{S1}$$

On the lefthand side Eq. S1, $P(N, n)$ is the probability of $N$ loops and $n$ cut loops, $P(n|N, p_{cut})$ is the probability that RAG cuts $n$ loops conditional on the $N$ total loops examined and the

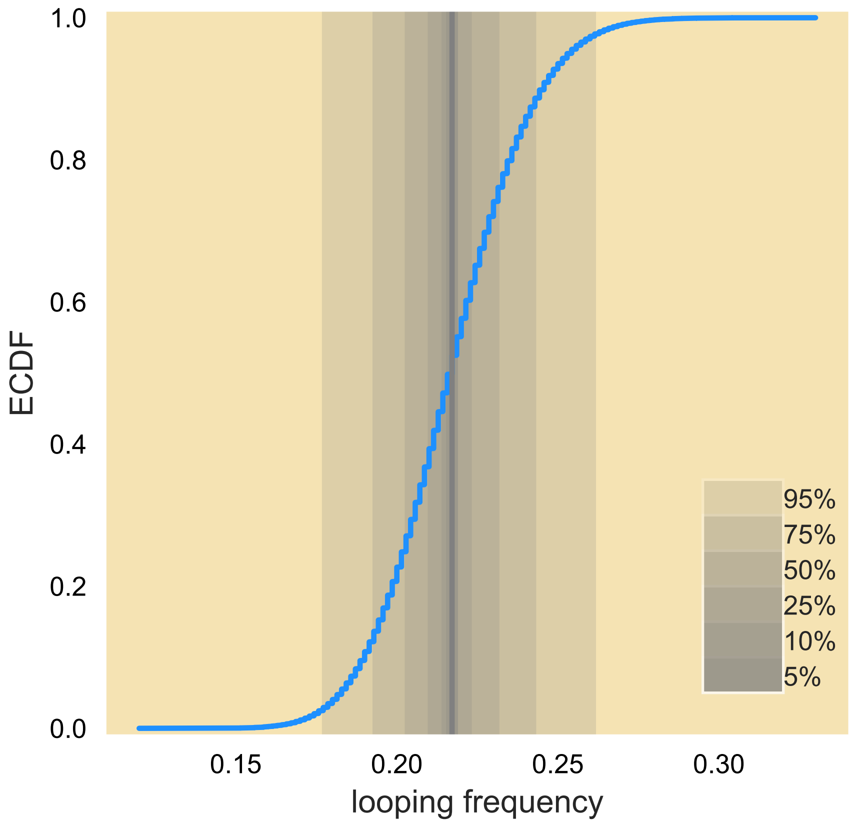

**Figure S2:** Bootstrapped looping frequency and confidence intervals for the V4-57-1 reference sequence. Empirical CDFs of the bootstrapped looping frequency with 5%, 10%, 25%, 50%, 75% and 95% confidence intervals as represented by the color bar.

probability that RAG cuts a given loop $p_{cut}$. $P(N, p_{cut})$ is the probability of getting $N$ total loops and that RAG has a cut probability $p_{cut}$ for the RSS. A rearrangement of the equation shows that

$$P(p_{cut}|N, n) = \frac{P(n|N, p_{cut})P(N, p_{cut})}{P(N, n)}. \tag{S2}$$

We can further simplify this equation by noting that the probability of getting $N$ loops and a cut probability $p_{cut}$ are independent values. This is evident from the fact that we could have carried out more TPM experiments and in principle $p_{cut}$ should not change from increasing the sample size of loops observed. Thus,

$$P(N, p_{cut}) = P(N)P(p_{cut}). \tag{S3}$$

Furthermore, we can further simplify the probability function in the denominator from noticing that the probability of having $N$ total loops and $n$ cut loops can be pieced apart as the probability of having $n$ cut loops given $N$ total loops times the probability of having $N$ total loops to begin with, or

$$P(N, n) = P(n|N)P(N). \tag{S4}$$

Inserting equations S3 and S4 into equation S2 gives us

$$P(p_{cut}|N, n) = \frac{P(n|N, p_{cut})P(N)P(p_{cut})}{P(n|N)P(N)},$$

$$= \frac{P(n|N, p_{cut})P(p_{cut})}{P(n|N)}. \quad \text{(S5)}$$

We wish to determine the conditional function on the left of Eq. S5, which we will term our posterior distribution. Here, we construct our posterior distribution from inputting the probabilities on the righthand side of the equation.

We first determine $P(n|N, p_{cut})$. This conditional probability function is the probability that we observe $n$ loops cut considering we observe $N$ loops forming and if the paired complex has a probability of cutting $p_{cut}$. Here, we would expect that this is similar to flipping a biased coin $N$ times and seeing how many instances heads comes up when the true value of the coin coming up heads is $p_{cut}$. In this case, we expect this conditional probability to be binomially distributed:

$$P(n|N, p_{cut}) = \frac{N!}{n!(N-n)!}(p_{cut})^n(1-p_{cut})^{N-n}. \quad \text{(S6)}$$

Next, we would like to determine $P(p_{cut})$. This is our prior distribution and, because this probability function is not conditioned on any data, this distribution function simply comes from our *a priori* knowledge of $p_{cut}$ independent of the data we have in hand. Here, we choose to say that the only knowledge we have of this parameter is that it, like all probabilities, is bounded between zero and one. We assume that $p_{cut}$ can take any value between zero and one equally. Thus,

$$P(p_{cut}) = \begin{cases} 1 & 0 \leq p_{cut} \leq 1, \\ 0 & \text{otherwise.} \end{cases} \quad \text{(S7)}$$

Finally, we need to determine the probability that $n$ loops cut given $N$ observed loops. This probability is only conditioned on $N$ and not $p_{cut}$, so we can say that $n$ can take on any integer value between 0 and $N$, inclusive. Thus, we have a discrete uniform distribution:

$$P(n|N) = \frac{1}{N+1}. \quad \text{(S8)}$$

By assembling equations S6, S7 and S8 into equation S5, we get that

$$P(p_{cut}|N, n) = \frac{(N+1)!}{n!(N-n)!}(p_{cut})^n(1-p_{cut})^{N-n}. \quad \text{(S9)}$$

With the posterior distribution in hand, we compute the most probable value of $p_{cut}$ by determining where the derivative of the posterior distribution with respect to $p_{cut}$ is 0. For ease of calculation, we will take the logarithm of the posterior distribution and derive with respect to $p_{cut}$:

$$\begin{aligned} ln[P(p_{cut}|N, n)] &= ln\Big[\frac{(N+1)!}{n!(N-n)!}\Big] + n\, ln(p_{cut}) + (N-n)\, ln(1-p_{cut}), \\ \frac{d\, ln[P(p_{cut}|N, n)]}{d\, p_{cut}}\Big|_{p^*_{cut}} &= \frac{n}{p^*_{cut}} - \frac{N-n}{1-p^*_{cut}} = 0. \end{aligned} \quad \text{(S10)}$$

Eq. S10 then tells us that

$$p^*_{cut} = \frac{n}{N}. \quad \text{(S11)}$$

To calculate the variance of $p_{cut}$, we make the assumption that $N \gg 1$ and look to center about the most probable value, $p^*_{cut}$. With this assumption, we will approximate the posterior distribution as a Gaussian distribution. In order to see this in action, we will define $x \equiv p - p^*_{cut}$. Then Eq. S12 becomes

$$P(p_{cut}|N, n) = \frac{(N+1)!}{n!(N-n)!}(p^*_{cut} + x)^n(1 - p^*_{cut} - x)^{N-n}. \quad \text{(S12)}$$

We also invoke the rule that $ln\, n! \approx n\, ln\, n - n + \frac{1}{2}\, ln[2\pi n]$. We can then determine the prefactor of the posterior distribution. Specifically,

$$\begin{aligned}
\frac{(N+1)!}{n!(N-n)!} &= exp\{ln[(N+1)!] - ln\, n! - ln[(N-n)!]\}, \\
&\approx exp\{(N+1)ln(N+1) - (N+1) + \frac{1}{2}ln[2\pi\,(N+1)] - n\, ln\, n + n - \frac{1}{2}ln(2\pi n) \\
&\qquad - (N-n)ln(N-n) + (N-n) - \frac{1}{2}ln[2\pi(N-n)]\}, \\
&\approx exp\Big\{(N+1)\Big[ln\, N + ln\Big(1+\frac{1}{N}\Big)\Big] - 1 - n\, ln\, n - (N-n)\Big[ln\, N + ln\Big(1-\frac{n}{N}\Big)\Big] \\
&\qquad + \frac{1}{2}ln\Big[\frac{N+1}{2\pi\, n(N-n)}\Big]\Big\}, \\
&\approx exp\Big\{(N+1)(\frac{1}{N} + \frac{1}{2N^2}) - 1 - n\, ln\, n + n\, ln\, N - (N-n)ln(1-p^*_{cut}) \\
&\qquad + \frac{1}{2}ln\Big[\frac{N^3}{2\pi\, n(N-n)}\Big]\Big\}, \\
&\approx \frac{1}{\sqrt{2\pi\frac{n(N-n)}{N^3}}} exp\Big\{-n\, ln\Big(p^*_{cut}\Big) - N(1-p^*_{cut})ln(1-p^*_{cut})\Big\}. \qquad \text{(S13)}
\end{aligned}$$

Here, we make simplifying assumptions, such as that $N+1 \approx N$ and Taylor expansions for $\frac{1}{N}$.

With the prefactor taken care of, we can rework the entire posterior distribution.

$$\begin{aligned}
P(p_{cut}|N,n) &\approx \frac{1}{\sqrt{2\pi\frac{n(N-n)}{N^3}}} exp\Big\{-n\, ln\Big(p^*_{cut}\Big) - N(1-p^*_{cut})ln(1-p^*_{cut}) + n\, ln(p^*_{cut}+x) \\
&\qquad + (N-n)ln(1-p^*_{cut}-x)\Big\}, \\
&\approx \frac{1}{\sqrt{2\pi\frac{n(N-n)}{N^3}}} exp\Big\{-n\, ln\Big(p^*_{cut}\Big) - N(1-p^*_{cut})ln(1-p^*_{cut}) + n\Big[ln(p^*_{cut}) + ln(1+\frac{x}{p^*_{cut}})\Big] \\
&\qquad + (N-n)\Big[ln(1-p^*_{cut}) + ln(1-\frac{x}{1-p^*_{cut}})\Big]\Big\}, \\
&\approx \frac{1}{\sqrt{2\pi\frac{n(N-n)}{N^3}}} exp\Big\{n\Big[ln(1+\frac{x}{p^*_{cut}})\Big] + (N-n)\Big[ln(1-\frac{x}{1-p^*_{cut}})\Big]\Big\}, \\
&\approx \frac{1}{\sqrt{2\pi\frac{n(N-n)}{N^3}}} exp\Big\{n\Big[\frac{x}{p^*_{cut}} - \frac{x^2}{2{p^*_{cut}}^2}\Big] + (N-n)\Big[-\frac{x}{1-p^*_{cut}} - \frac{x^2}{2(1-p^2_{cut})^2}\Big]\Big\}, \\
&\approx \frac{1}{\sqrt{2\pi\frac{n(N-n)}{N^3}}} exp\Big\{N\, x - n\frac{x^2}{2{p^*_{cut}}^2} - N\, x - (N-n)\frac{x^2}{2(1-p^*_{cut})^2}\Big\}, \\
&\approx \frac{1}{\sqrt{2\pi\frac{n(N-n)}{N^3}}} exp\Big\{-n\frac{x^2}{2{p^*_{cut}}^2} - (N-n)\frac{x^2}{2(1-p^*_{cut})^2}\Big\}, \\
&\approx \frac{1}{\sqrt{2\pi\frac{n(N-n)}{N^3}}} exp\Big\{-N\frac{x^2}{2p^*_{cut}} - N\frac{x^2}{2(1-p^*_{cut})}\Big\}, \\
&\approx \frac{1}{\sqrt{2\pi\frac{n(N-n)}{N^3}}} exp\Big\{-\frac{N\, x^2}{2}\Big(\frac{1}{p^*_{cut}} + \frac{1}{1-p^*_{cut}}\Big)\Big\},
\end{aligned}$$

$$\approx \frac{1}{\sqrt{2\pi \frac{n(N-n)}{N^3}}} exp\Big\{ -\frac{N\,x^2}{2}\Big(\frac{1}{p^*_{cut}(1-p^*_{cut})}\Big)\Big\},$$

$$\approx \frac{1}{\sqrt{2\pi \frac{n(N-n)}{N^3}}} exp\Big\{ -\frac{(p-p^*_{cut})^2}{2\Big[\frac{n\,(N-n)}{N^3}\Big]}\Big\}. \tag{S14}$$

Eq. S14 tells us that, not only is this Gaussian approximation centered at the most probable value of $p_{cut} = p^*_{cut}$, as we would expect, but also that the distribution has a variance of $\sigma^2 = \frac{n(N-n)}{N^3}$. Thus, we report $p^*_{cut} = \frac{n}{N}$ and $\sigma^2 = \frac{n(N-n)}{N^3}$ in Fig. 3C and 4C of the main text.

#### S2.4 Modeling Exit from the Paired Complex As A Poisson Process

As discussed in the main text, we attempted to model the kinetics of unlooping and exiting of the paired complex state. In the case of exit, we considered that every paired complex had one of two fates; either the DNA was cleaved and the observed tethered bead was lost or the paired complex dissociated, releasing the bead to its full-length tethered state. We consider these two fates as independent yet competing processes. Under the independence assumption, we can model each process individually as a Poisson process where the time of leaving the paired complex (either through cleavage or unlooping) is exponentially distributed. Mathematically, we can state that the probability of leaving the paired complex at time $t_{\text{leave}}$ is defined as

$$P(t_{\text{leave}}\,|\,k_{\text{leave}}) = k_{\text{leave}} e^{-k_{\text{leave}} t_{\text{leave}}}, \tag{S15}$$

where the leaving rate $k_{\text{leave}}$ is defined as the sum of the two independent rates,

$$k_{\text{leave}} = k_{\text{cut}} + k_{\text{unloop}}. \tag{S16}$$

Therefore, given a collection of paired complex dwell times $t_{\text{leave}}$, we can estimate the most-likely value for $k_{\text{leave}}$ providing insight on whether exiting the paired complex can be modeled as a Poisson process.

Rather than reporting a single value, we can determine the probability distribution of the parameter $k_{\text{leave}}$. This distribution, termed the posterior distribution, can be computed by Bayes' theorem as

$$P(k_{\text{leave}}\,|\,t_{\text{leave}}) = \frac{P(t_{\text{leave}}\,|\,k_{\text{leave}})P(k_{\text{leave}})}{P(t_{\text{leave}})}. \tag{S17}$$

The posterior distribution $P(k_{\text{leave}}\,|\,t_{\text{leave}})$ defines the probability of a leaving rate *given* a set of measurements $t_{\text{leave}}$. This distribution is dependent on the likelihood of observing the dwell time distribution *given* a leaving rate, represented by $P(t_{\text{leave}}\,|\,k_{\text{leave}})$. All prior information we have about what the leaving rate *could* be is captured by $P(k_{\text{leave}})$ which is entirely independent of the data. The denominator in Eq. S17 defines the probability distribution of the data marginalized over all values of the leaving rate. For our purposes, this term serves as a normalization constant and will be neglected.

We are now tasked with defining functional forms for the various probabilities enumerated in Eq. S17. The likelihood already matches the definition in Eq. S15, so we assign our likelihood as a simple exponential distribution parameterized by the leaving rate. Choosing a functional form for the prior distribution $P(k_{\text{leave}})$ is a much more subjective process. As such, we outline our thinking below.

As written in Eq. S15, $k_{\text{leave}}$ has dimensions of inverse time, meaning that particularly long-lived paired complexes will have $k_{\text{leave}} < 1$ whereas a sequence with unstable paired complexes will have $k_{\mathrm{leave}} > 1$. As we remain ignorant of our data, we consider both of these extremes to be valid values for the leaving rate. However, this parameterization raises technical issues with estimating $k_{\mathrm{leave}}$ computationally. We sample the complete posterior using Markov chain Monte Carlo, a computational technique in which the posterior is explored via a biased random walk depending on the gradient of the local probability landscape. With such a widely constrained parameter, effectively sampling very small values of $k_{\mathrm{leave}}$ becomes more difficult than larger values. We can avoid this issue by reparameterizing Eq. S15 in terms of the inverse leaving rate $\tau_{\mathrm{leave}} = \frac{1}{k_{\mathrm{leave}}}$ so that

$$P(t_{\mathrm{leave}} \,|\, \tau_{\mathrm{leave}}) = \frac{1}{\tau_{\mathrm{leave}}} e^{t_{\mathrm{leave}}/\tau_{\mathrm{leave}}}. \tag{S18}$$

Our parameter of interest now has dimensions of time and can be interpreted as the average life time of a paired complex or, more precisely, the waiting time for the arrival of a Poisson process.

While it is tempting to default to a completely uninformative prior for $\tau_{\mathrm{leave}}$ to avoid introducing any bias into our parameter estimation, we do have some intuition for what the bounds of the value could be. For example, it is mathematically impossible for the leaving rate to be less than zero. We can also find it unlikely that the leaving rate is *exactly* zero as that would imply irreversible formation of the paired complex. We can therefore say that the value for the leaving rate is positive and can asymptotically approach zero. As we have designed the experiment to actually observe the entry and exit of the paired complex state, we can set a soft upper bound for the leaving rate to be the length of our typical experiment, 60 minutes. With these bounds in place, we can assign some probability distribution between them where it is impossible to equal zero and improbable but not impossible to exceed 60 minutes.

A good choice for such a distribution is an inverse Gamma distribution which has the form

$$P(\tau_{\mathrm{leave}} \,|\, \alpha, \beta) = \frac{1}{\Gamma(\alpha)} \frac{\beta^{\alpha}}{\tau_{\mathrm{leave}}^{(\alpha+1)}} e^{-\beta/\tau_{\mathrm{leave}}}, \tag{S19}$$

where $\alpha$ and $\beta$ correspond to the number of arrivals of a Poisson process and their rate of arrival, respectively. Given that only one arrival is necessary to exit a paired complex, we choose $\alpha$ to be approximately equal to 1 and $\beta$ to be approximately 10. This meets our conditions described previously of asymptotically approaching zero and rarely exceeding 60 minutes.

Combining Eq. S18 and Eq. S19 yields the complete posterior distribution. We sampled this distribution for each RSS independently using Markov chain Monte Carlo. Specifically, we used Hamiltonian Markov chain Monte Carlo as is implemented in the Stan probabilistic programming language [6]. The code used to sample this distribution can be accessed on the paper website or GitHub repository.

### S3 Posterior Distributions of the Endogenous Sequences

Fig. S3 gives the full posterior distributions of the cutting probability for each of the endogenous RSSs. We see clearly that between the two RSSs flanking the DFL16.1 gene segment that RAG is more successful at cleaving the RSS on the 3' side of the gene segment than the RSS on the 5' end. In examining the RSSs adjacent to endogenous V$\kappa$ gene segments, we see that the cutting probability is not differentiable across most of the RSSs, but cleavage is dramatically reduced when RAG interacts with the V5-43, V8-18 and V6-15 RSSs. We find that the number of paired complexes formed with the V8-18 12RSS is low to begin with, leading to an uninformative posterior distribution, whereas the V6-15 12RSS may suffer a low cleavage probability due to the T immediately adjacent to the heptamer in the coding flank region, which has been known to broadly reduce recombination efficiency [7, 8, 9].

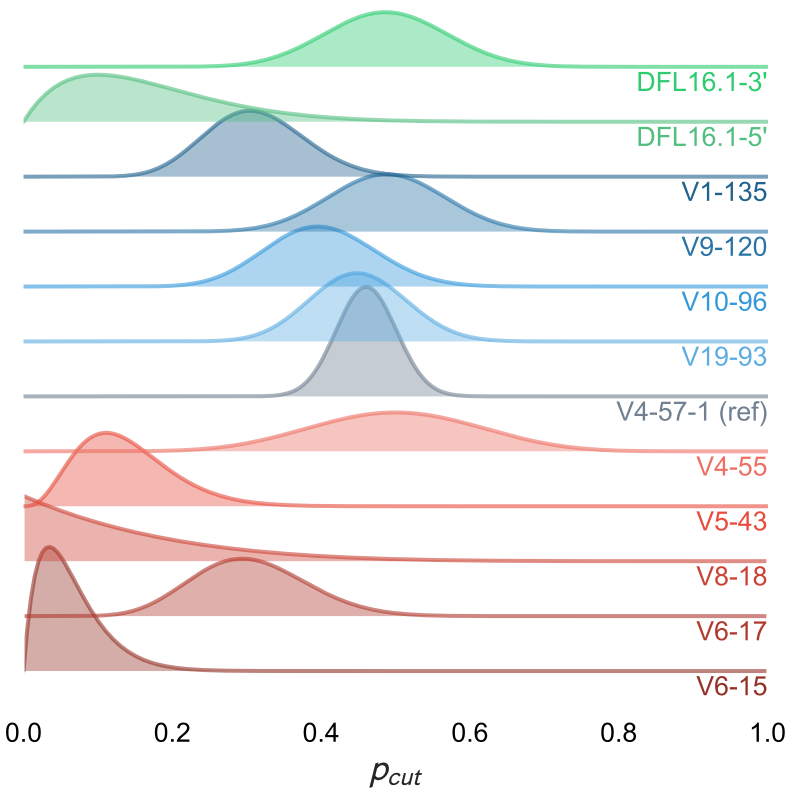

**Figure S3:** Posterior distributions of the cutting probabilities as derived in SI Section S2.3 for the endogenous 12RSSs studied. The top-to-bottom order of the endogenous RSSs is the same as their left-to-right ordering in Fig. 3. Height of the distribution is proportional to the probability of the $p_{\rm cut}$ value.

#### S4 Coding Flank Contributions

For our study of the endogenous RSSs, we also modified the coding flanks adjacent to the RSSs during the cloning process to construct the DNA tethers. As shown in table S1, most of these coding flanks have A and C nucleotides in the two or three base pairs upstream of the heptamer region. However, recent structural work have shown direct contacts between RAG1 residues and the coding flank [10, 11, 12]. Furthermore, various bulk assays have demonstrated that coding flank sequence can affect recombination efficiency [7, 8, 9]. These bulk assays suggest that coding flanks with A and C nucleotides near the heptamer tend to recombine more efficiently than those that have Ts instead. In attempting to determine whether these A- and C-rich coding flanks have much of an influence on the RAG-RSS dynamics, we looked to two pairs of TPM constructs where within each pair the RSS is identical, but the coding flank sequence is different.

Fig. S4 shows TPM results on the V4-57-1, or reference, RSS and a single bp change at the nucleotide immediately adjacent to the heptamer, where there is a C-to-A alteration. We find here no distinguishable difference in looping frequency or cleavage probability. Furthermore, we find that the dwell time distributions for PCs that cut, PCs that unloop, and both are identical between the reference and altered coding flank. This finding suggests that at least a single change from C to A near the heptamer does not have a dramatic effect on the RAG-RSS interaction.

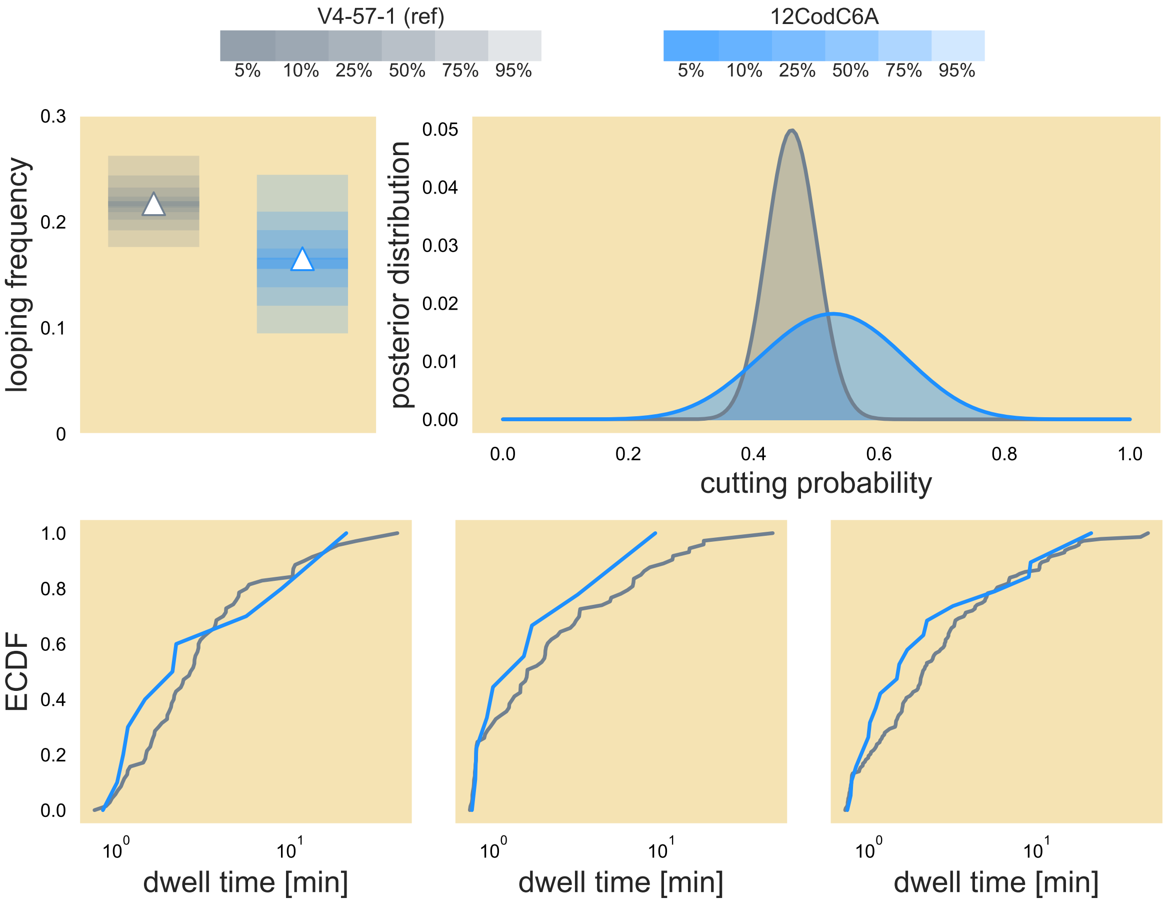

**Figure S4:** V4-57-1 (reference) RSS (grey) and coding flank change (blue) comparison of looping frequency, posterior distribution of the cutting probability and ECDFs of PC lifetimes for PCs that cut, those that unloop, and both.

We also examined two coding flanks that differ by multiple base pairs. The V4-55 RSS differs from the reference sequence at the first position of the spacer, where the C for the reference is changed to an A for the V4-55 RSS. However, the coding flank sequence differs at five nucleotides. Furthermore, the 6-bp coding flank of V4-55 is composed entirely of Cs and As and removes the Gs and Ts on the reference sequence at the first, third, and fourth positions of the coding flank (where we index one as six base pairs from the start of the heptamer and six as immediately adjacent). We thus compared the C-to-A change at the spacer position 1 on the reference sequence with the V4-55 coding flank. As Fig. S5 illustrates that despite the significant difference in sequence between these two constructs at the coding flank, our TPM assay reports little difference that separates these two sequences in looping frequency, dwell time distributions or cutting probability. We thus find that for most of the endogenous RSSs studied, the coding flank has little effect on the overall RAG-RSS interaction. This does not rule out the possibility that Gs or Ts in the first three positions immediately adjacent to the RSS can alter the RAG-RSS dynamics.

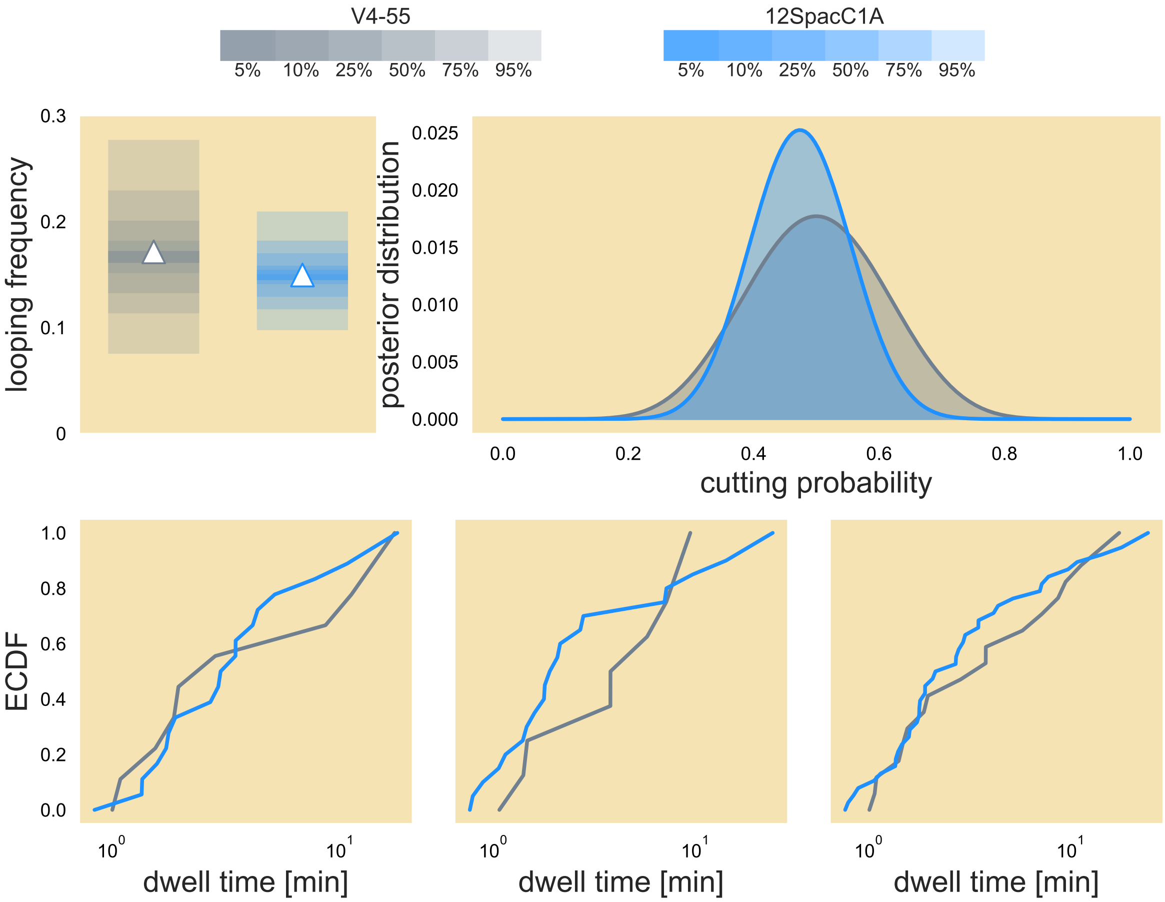

**Figure S5:** V4-55 12RSS (grey) and C-to-A change at spacer position 1 (blue) comparison of looping frequency, posterior distribution of the cutting probability and ECDFs of PC lifetimes for PCs that cut, those that unloop, and both

#### S5 $Ca^{2+}$-$Mg^{2+}$ Looping Frequency Comparisons

Although we directly compared the dwell time distributions of three RSS constructs between when the RAG reaction buffer contained $Mg^{2+}$ to allow for nicking and buffer containing $Ca^{2+}$ to prevent nicking, we wanted to know whether the looping frequency would increase when RAG is prohibited from nicking. Our intuition comes from recognizing that without the ability to cleave the DNA, RAG can only release one of the RSSs and leave the paired complex state without cutting the DNA tether. As a result, RAG has an opportunity to form the paired complex with the same DNA tether. We expect that the looping frequency should either increase or remain the same in the $Ca^{2+}$ environment as compared to when $Mg^{2+}$ is used. Fig. S6 shows that these two outcomes result. Fig. S6A and S6C show that RAG forms the paired complex more frequently with the reference sequence and the G-to-T change at the eleventh position of the reference spacer sequence when the reaction occurs in $Ca^{2+}$. Furthermore, we see that undergoing the reaction with the A-to-T alteration at heptamer position four in $Ca^{2+}$ does not induce much change in the looping frequency as compared to a $Mg^{2+}$ environment (Fig. S6). Of interest is the fact that the spacer variant, which has a slightly larger measured looping frequency than the reference sequence in $Mg^{2+}$ with overlapping 95% confidence intervals, clearly undergoes a more dramatic increase in looping frequency than the reference sequence when the salt is $Ca^{2+}$. This observation shows that PC formation is more favorable for the spacer variant than the reference sequence. Observed holistically, we find that RAG in the absence of nicking can form loops at least as frequently as when it when it can nick the DNA.

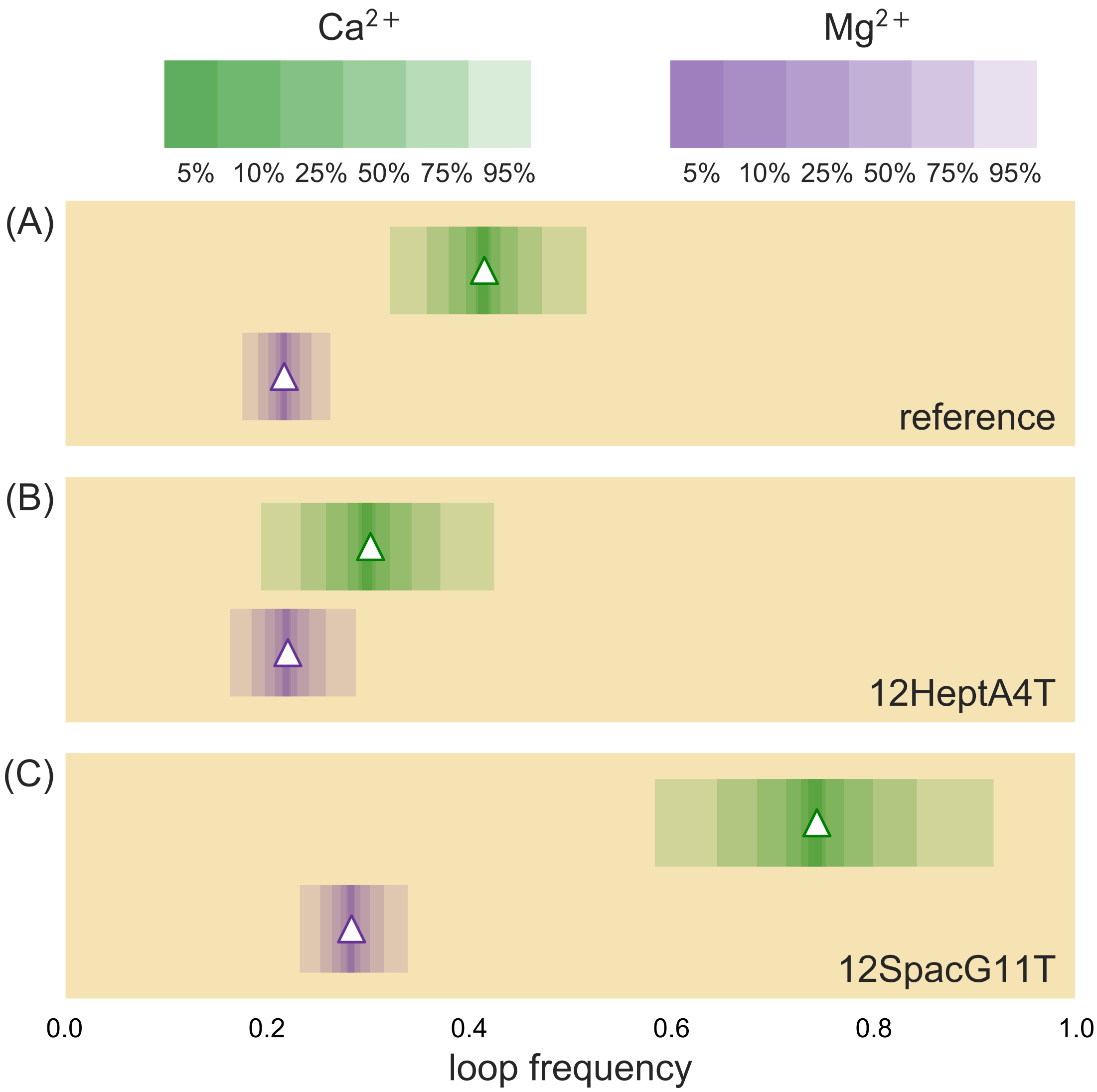

**Figure S6:** $Ca^{2+}$ (green) and $Mg^{2+}$ (purple) looping frequencies for (A) reference 12RSS, (B) A-to-T change at the fourth position of the heptamer and (C) G-to-T change at the eleventh position of the spacer. Measured looping frequency shown as the triangles. Going from darker shading to lighter shading in rectangle bar indicates increasing of confidence interval percentage of the looping frequency from the bootstrapping method discussed in section S2.2.

### S6 Endogenous RSS Sequences

| Endogenous 12RSS | Sequence |
|---|---|
| DFL16.1-3' | AGCTAC CACAGTGCTATATCCATCAGCAAAAACC |
| DFL16.1-5' | AATAAA CACAGTAGTAGATCCCTTCACAAAAAGC |
| V1-135 | TCCTCA CACAGTGATTCAGACCCGAACAAAAACT |
| V9-120 | TCCTCC CACAGTGATACAAATCATAACATAAACC |
| V10-96 | TCCTCC CACAATGATATAAGTCATAACATAAACC |
| V19-93 | TCTACC CACAGTGATACAAATCATAACAAAAACC |
| V4-57-1 (reference) | GTCGAC CACAGTGCTACAGACTGGAACAAAAACC |
| V4-55 | CACCCA CACAGTGATACAGACTGGAACAAAAACC |
| V5-43 | GCCTCA CACAGTGATGCAGACCATAGCAAAAATC |
| V8-18 | TCCCCC CACAGAGCTTCAGCTGCCTACACAAACC |
| V6-17 | TCCTCC CACAGTGCTTCAGCCTCCTACACAAACC |
| V6-15 | TCCTCT CACAGTACTTCAGCCTCCTACATAAACC |
| **J$\kappa$1 23RSS** | GGATCC CACAGTGGTAGTACTCCACTGTCTGGCTGTACAAAAACC |

Table S1: **Table of endogenous 12RSS sequences.** The 6 base pairs before the heptamer, known as the coding flank, is changed in the endogenous RSS studies and is included here. The spacer sequence for each RSS is underlined. Bold blue letters in the heptamer and nonamer regions denote deviations from the consensus sequences, CACAGTG and ACAAAAACC, respectively. The bottom sequence is of the J$\kappa$1 23RSS applied in all of the DNA constructs used in TPM.

### S7 Cloning a Different 12RSS in Plasmids

To generate the synthetic RSSs used in this work, we used overhang PCR (polymerase chain reaction) and subsequently Gibson assembly (NEB Biolabs) to generate plasmids with the desired change. We selected the endogenous sequence V4-57-1 to serve as the "reference" sequence from which all synthetic RSSs were made. This sequence has been used previously [2] and exhibits a reasonable dwell time distribution, has moderately high looping frequency (compared to the other endogenous sequences), and has close to a 50% cleavage probability, as is shown in this study. This 12RSS sequence is located within the a pZE12 plasmid backbone [13]. The new RSS were inserted into this plasmid via overhang PCR with forward and reverse oligonucleotide primers (IDT) that contain a 15 base-pair overlap with the desired alteration in the middle of the sequence. The primers used in this work are listed in tables S2 and S3.

After purification of the PCR fragment and DpnI digestion (NEB Biolabs) of the PCR template, the fragment was circularized using Gibson assembly [14] and transformed into DH5$\alpha$ *Escherichia coli.* Transformants were then cultured and stored for plasmid purification and sequence verification.

### S8 Synthetic 12RSS Primers

Tables S2 and S3 gives the list of primers used to construct the synthetic and endogenous RSSs. For synthetic RSSs, we apply the nomenclature '12' to denote that the 12RSS is altered, the region of the RSS where the change is made ('Hept' = heptamer, 'Non' = nonamer, 'Spac' = spacer, 'Cod' = coding flank), the original nucleotide, the position number in the region, where indexing starts at 1 and finally the new nucleotide. Therefore, if a change is made to the eighth position of the spacer, where a C is altered to T, the RSS is denoted '12SpacC8T'.

| Synthetic 12RSS | Primer |
|---|---|
| 12CodC6A (Fwd) | AACACAGTGCTACAGACTGGAACAAAAACCCTGCAGTC |
| 12CodC6A (Rev) | CTGTAGCACTGTGTTCGACCTGCAGCCCAAGCG |
| 12HeptC3G (Fwd) | ACCAGAGTGCTACAGACTGGAACAAAAACCCTGCAGTC |
| 12HeptC3G (Rev) | CTGTAGCACTCTGGTCGACCTGCAGCCCAAGCG |
| 12HeptC3T (Fwd) | ACCATAGTGCTACAGACTGGAACAAAAACCCTGCAGTC |
| 12HeptC3T (Rev) | CTGTAGCACTATGGTCGACCTGCAGCCCAAGCG |
| 12HeptA4T (Fwd) | ACCACTGTGCTACAGACTGGAACAAAAACCCTGCAGTC |
| 12HeptA4T (Rev) | CTGTAGCACAGTGGTCGACCTGCAGCCCAAGCG |
| 12HeptG5A (Fwd) | ACCACAATGCTACAGACTGGAACAAAAACCCTGCAGTC |
| 12HeptG5A (Rev) | CTGTAGCATTGTGGTCGACCTGCAGCCCAAGCG |
| 12HeptG5C (Fwd) | ACCACACTGCTACAGACTGGAACAAAAACCCTGCAGTC |
| 12HeptG5C (Rev) | CTGTAGCAGTGTGGTCGACCTGCAGCCCAAGCG |
| 12HeptT6A (Fwd) | ACCACAGAGCTACAGACTGGAACAAAAACCCTGCAGTC |
| 12HeptT6A (Rev) | CTGTAGCTCTGTGGTCGACCTGCAGCCCAAGCG |
| 12HeptT6C (Fwd) | ACCACAGCGCTACAGACTGGAACAAAAACCCTGCAGTC |
| 12HeptT6C (Rev) | CTGTAGCGCTGTGGTCGACCTGCAGCCCAAGCG |
| 12HeptG7A (Fwd) | ACCACAGTACTACAGACTGGAACAAAAACCCTGCAGTC |
| 12HeptG7A (Rev) | CTGTAGTACTGTGGTCGACCTGCAGCCCAAGCG |
| 12HeptG7C (Fwd) | ACCACAGTCCTACAGACTGGAACAAAAACCCTGCAGTC |
| 12HeptG7C (Rev) | CTGTAGGACTGTGGTCGACCTGCAGCCCAAGCG |
| 12HeptG7T (Fwd) | ACCACAGTTCTACAGACTGGAACAAAAACCCTGCAGTC |
| 12HeptG7T (Rev) | CTGTAGAACTGTGGTCGACCTGCAGCCCAAGCG |
| 12SpacC1A (Fwd) | ACCACAGTGATACAGACTGGAACAAAAACCCTGCAGTC |
| 12SpacC1A (Rev) | CTGTATCACTGTGGTCGACCTGCAGCCCAAGCG |
| 12SpacC1G (Fwd) | ACCACAGTGGTACAGACTGGAACAAAAACCCTGCAGTC |
| 12SpacC1G (Rev) | CTGTACCACTGTGGTCGACCTGCAGCCCAAGCG |
| 12SpacA3G (Fwd) | ACCACAGTGCTGCAGACTGGAACAAAAACCCTGCAGTC |
| 12SpacA3G (Rev) | CTGCAGCACTGTGGTCGACCTGCAGCCCAAGCG |
| 12SpacA3T (Fwd) | ACCACAGTGCTTCAGACTGGAACAAAAACCCTGCAGTC |
| 12SpacA3T (Rev) | CTGAAGCACTGTGGTCGACCTGCAGCCCAAGCG |
| 12SpacC4G (Fwd) | ACCACAGTGCTAGAGACTGGAACAAAAACCCTGCAGTC |
| 12SpacC4G (Rev) | CTCTAGCACTGTGGTCGACCTGCAGCCCAAGCG |
| 12SpacC4T (Fwd) | ACCACAGTGCTATAGACTGGAACAAAAACCCTGCAGTC |
| 12SpacC4T (Rev) | CTATAGCACTGTGGTCGACCTGCAGCCCAAGCG |
| 12SpacG6A (Fwd) | ACCACAGTGCTACAAACTGGAACAAAAACCCTGCAGTC |
| 12SpacG6A (Rev) | TTGTAGCACTGTGGTCGACCTGCAGCCCAAGCG |
| 12SpacG6T (Fwd) | ACCACAGTGCTACATACTGGAACAAAAACCCTGCAGTC |
| 12SpacG6T (Rev) | ATGTAGCACTGTGGTCGACCTGCAGCCCAAGCG |
| 12SpacA7C (Fwd) | ACCACAGTGCTACAGCCTGGAACAAAAACCCTGCAGTC |
| 12SpacA7C (Rev) | TTGTAGCACTGTGGTCGACCTGCAGCCCAAGCG |
| 12SpacA7G (Fwd) | ACCACAGTGCTACAGGCTGGAACAAAAACCCTGCAGTC |
| 12SpacA7G (Rev) | TTGTAGCACTGTGGTCGACCTGCAGCCCAAGCG |
| 12SpacC8T (Fwd) | ACCACAGTGCTACAGATTGGAACAAAAACCCTGCAGTC |

| | |
|---|---|
| 12SpacC8T (Rev) | TTGTAGCACTGTGGTCGACCTGCAGCCCAAGCG |
| 12SpacT9A (Fwd) | ACCACAGTGCTACAGAC**A**GGAACAAAAACCCTGCAGTC |
| 12SPacT9A (Rev) | TTGTAGCACTGTGGTCGACCTGCAGCCCAAGCG |
| 12SpacT9C (Fwd) | ACCACAGTGCTACAGAC**C**GGAACAAAAACCCTGCAGTC |
| 12SpacT9C (Rev) | TTGTAGCACTGTGGTCGACCTGCAGCCCAAGCG |
| 12SpacT9G (Fwd) | ACCACAGTGCTACAGAC**G**GGAACAAAAACCCTGCAGTC |
| 12SpacT9G (Rev) | TTGTAGCACTGTGGTCGACCTGCAGCCCAAGCG |
| 12SpacG10A (Fwd) | ACCACAGTGCTACAGACT**A**GAACAAAAACCCTGCAGTC |
| 12SpacG10A (Rev) | CTGTAGCACTGTGGTCGACCTGCAGCCCAAGCG |
| 12SpacG10C (Fwd) | ACCACAGTGCTACAGACT**C**GAACAAAAACCCTGCAGTC |
| 12SpacG10C (Rev) | CTGTAGCACTGTGGTCGACCTGCAGCCCAAGCG |
| 12SpacG10T (Fwd) | ACCACAGTGCTACAGACT**T**GAACAAAAACCCTGCAGTC |
| 12SpacG10T (Rev) | CTGTAGCACTGTGGTCGACCTGCAGCCCAAGCG |
| 12SpacG11A (Fwd) | ACCACAGTGCTACAGACTG**A**AACAAAAACCCTGCAGTC |
| 12SpacG11A (Rev) | CTGTAGCACTGTGGTCGACCTGCAGCCCAAGCG |
| 12SpacG11C (Fwd) | ACCACAGTGCTACAGACTG**C**AACAAAAACCCTGCAGTC |
| 12SpacG11C (Rev) | CTGTAGCACTGTGGTCGACCTGCAGCCCAAGCG |
| 12SpacG11T (Fwd) | ACCACAGTGCTACAGACTG**T**AACAAAAACCCTGCAGTC |
| 12SpacG11T (Rev) | CTGTAGCACTGTGGTCGACCTGCAGCCCAAGCG |
| 12SpacA12C (Fwd) | ACCACAGTGCTACAGACTGG**C**ACAAAAACCCTGCAGTC |
| 12SpacA12C (Rev) | CTGTAGCACTGTGGTCGACCTGCAGCCCAAGCG |
| 12SpacA12T (Fwd) | ACCACAGTGCTACAGACTGG**T**ACAAAAACCCTGCAGTC |
| 12SpacA12T (Rev) | CTGTAGCACTGTGGTCGACCTGCAGCCCAAGCG |
| 12NonA1G (Fwd) | ACCACAGTGCTACAGACTGGA**G**CAAAAACCCTGCAGTC |
| 12NonA1G (Rev) | CTGTAGCACTGTGGTCGACCTGCAGCCCAAGCG |
| 12NonA3C (Fwd) | ACCACAGTGCTACAGACTGGAAC**C**AAAACCCTGCAGTC |
| 12NonA3C (Rev) | CTGTAGCACTGTGGTCGACCTGCAGCCCAAGCG |
| 12NonA4C (Fwd) | ACCACAGTGCTACAGACTGGAACA**C**AAACCCTGCAGTC |
| 12NonA4C (Rev) | CTGTAGCACTGTGGTCGACCTGCAGCCCAAGCG |
| 12NonA4T (Fwd) | ACCACAGTGCTACAGACTGGAACA**T**AAACCCTGCAGTC |
| 12NonA4T (Rev) | CTGTAGCACTGTGGTCGACCTGCAGCCCAAGCG |
| 12NonA5T (Fwd) | ACCACAGTGCTACAGACTGGAACAA**T**AACCCTGCAGTC |
| 12NonA5T (Rev) | CTGTAGCACTGTGGTCGACCTGCAGCCCAAGCG |
| 12NonC8G (Fwd) | GCTACAGACTGGAACAAAAA**G**CCTGCAGTCAACCTCGA |
| 12NonC8G (Rev) | TTTGTTCCAGTCTGTAGCACTGTGGTCGACCTGCAG |
| 12NonC8T (Fwd) | GCTACAGACTGGAACAAAAA**T**CCTGCAGTCAACCTCGA |
| 12NonC8T (Rev) | TTTGTTCCAGTCTGTAGCACTGTGGTCGACCTGCAG |
| 12NonC9T (Fwd) | GCTACAGACTGGAACAAAAAC**T**CTGCAGTCAACCTCGA |
| 12NonC9T (Rev) | TTTGTTCCAGTCTGTAGCACTGTGGTCGACCTGCAG |

Table S2: Forward (Fwd) and reverse (Rev) primers of synthetic RSSs. Underlined sequence denotes the region where change is made. Bold-faced letter denotes the new nucleotide.

| Endogenous 12RSS | Primer |
|---|---|
| DFL16.1-3' (Fwd) | AGCTACCACAGTGCTATATCCATCAGCAAAAACCCTGCAGTCGAGTAATGCA |
| DFL16.1-3' (Rev) | GGTTTTTGCTGATGGATATAGCACTGTGGTATTCGAAGCTTGAGCTCG |
| DFL16.1-5' (Fwd) | AATAAACACAGTAGTAGATCCCTTCACAAAAAGCCTGCAGTCGAGTAATGCA |
| DFL16.1-5' (Rev) | GCTTTTTGTGAAGGGATCTACTACTGTGGTATTCGAAGCTTGAGCTCG |
| V1-135 (Fwd) | TCCTCACACAGTGATTCAGACCCGAACAAAAACTCTGCAGTCAACCTCGAGAAACG |
| V1-135 (Rev) | AGTTTTTGTTCGGGTCTGAATCACTGTGTGAGGACTGCAGCCCAAGCGTGTAG |
| V9-120 (Fwd) | TCCTCCCACAGTGATACAAATCATAACATAAACCCTGCAGTCAACCTCGAGAAACG |
| V9-120 (Rev) | GGTTTATGTTATGATTTGTATCACTGTGGGAGGACTGCAGCCCAAGCGTGTAG |
| V10-96 (Fwd) | TCCTCCCACAATGATATAAGTCATAACATAAACCCTGCAGTCAACCTCGAGAAACG |
| V10-96 (Rev) | GGTTTATGTTATGACTTATATCATTGTGGGAGGACTGCAGCCCAAGCGTGTAG |
| V19-93 (Fwd) | TCTACCCACAGTGATACAAATCATAACAAAAACCCTGCAGTCAACCTCGAGAAACG |
| V10-93 (Rev) | GGTTTTTGTTATGATTTGTATCACTGTGGGTAGACTGCAGCCCAAGCGTGTAG |
| V4-55 (Fwd) | CACCCACACAGTGATACAGACTGGAACAAAAACCCTGCAGTCAACCTCGAGAAACG |
| V4-55 (Rev) | GGTTTTTGTTCCAGTCTGTATCACTGTGTGGGTGCTGCAGCCCAAGCGTGTAG |
| V5-43 (Fwd) | GCCTCACACAGTGATGCAGACCATAGCAAAAATCCTGCAGTCAACCTCGAGAAACG |
| V5-43 (Rev) | GATTTTTGCTATGGTCTGCATCACTGTGTGAGGCCTGCAGCCCAAGCGTGTAG |
| V8-18 (Fwd) | TCCCCCCACAGAGCTTCAGCTGCCTACACAAACCCTGCAGTCAACCTCGAGAAACG |
| V8-18 (Rev) | GGTTTGTGTAGGCAGCTGAAGCTCTGTGGGGGGACTGCAGCCCAAGCGTGTAG |
| V6-17 (Fwd) | TCCTCCCACAGTGCTTCAGCCTCCTACACAAACCCTGCAGTCAACCTCGAGAAACG |
| V6-17 (Rev) | GGTTTGTGTAGGAGGCTGAAGCACTGTGGGAGGACTGCAGCCCAAGCGTGTAG |
| V6-15 (Fwd) | TCCTCTCACAGTACTTCAGCCTCCTACATAAACCCTGCAGTCAACCTCGAGAAACG |
| V6-15 (Rev) | GGTTTATGTAGGAGGCTGAAGTACTGTGAGAGGACTGCAGCCCAAGCGTGTAG |

Table S3: Forward (Fwd) and reverse (Rev) primers for designing TPM constructs with endogenous 12RSSs. Underlined regions denote the heptamer and nonamer regions.

### S9 Protein Purification

#### S9.1 Murine core RAG1 and core RAG2 Co-Purification

Maltose-binding protein(MBP)-tagged murine core RAG1 and core RAG2 are co-transfected into HEK293-6E suspension cells using BioT transfection agent and are expressed in the cells for 48 hours. Cells are centrifuged and collected before resuspending with a lysis buffer consisting of cOmplete Ultra protease inhibitor and Tween-20 detergent before lysis through a cell disruptor. Lysate is centrifuged to remove the cell membrane and the supernatant containing expressed RAG is mixed with amylose resin to bind the MBP region to the resin before loading onto a chromatography gravity column. Amylose-attached RAG is then washed using lysis buffer, wash buffer containing salts before eluting with buffer containing high concentrations of maltose to out-compete the MBP on the resin. RAG-contained eluate is then concentrated and dialyzed in buffer containing 25 mM Tris-HCl (pH 8.0), 150 mM KCl, 2 mM DTT and 10% glycerol before snap-freezing 5-10 µL aliquots and storing at -80°C.

#### S9.2 HMGB1 Purification

Though not discussed extensively in this paper, the high mobility group box 1 (HMGB1) protein binds nonspecifically to DNA and helps facilitate RAG binding onto the RSS. A plasmid containing a His-tagged HMGB1 gene is transformed into BL21(DE3) cells and grown in liquid cultures until they reach an OD600 of 0.7. Cultures are then induced with isopropyl-$\beta$-D-1-thiogalactopyranoside (IPTG) to express HMGB1 for 4 hours at 30°C before cells are collected from the media. Cells are resuspended in binding buffer media containing cOmplete Ultra protease inhibitor, benzonase, Tween-20 and a low imidazole concentration before lysis through the cell disruptor. Lysate is cleared of membrane with an ultracentrifuge and loaded onto a nickel-NTA column to bind HMGB1. Nickel-bound HMGB1 is then washed with more binding buffer before washing with buffer containing low imidazole concentration. Washed HMGB1 are then eluted through the column with elution buffer containing higher concentration imidazole. Degraded HMGB1 is then removed by loading HMGB1 eluate onto SP column and collecting flow-through in 200 µL aliquots with an incrementally increasing salt gradient on the AKTA. Fractions that show highest change in voltage reading on the AKTA are run on a Western blot to confirm that protein of the correct size is collected before collecting. HMGB1 are transferred to a dialysis buffer containing 25 mM Tris-HCl (pH 8.0), 150 mM KCl, 2 mM DTT and 10% glycerol through a buffer-exchange centrifuge column before snap-freezing 5-10 µL aliquots and freezing at -80°C.
